# Advances in Protein Function Prediction from the Fifth CAFA Challenge

**DOI:** 10.64898/2026.04.27.716980

**Authors:** M. Clara De Paolis Kaluza, Rashika Ramola, Parnal Joshi, Damiano Piovesan, Walter Reade, Sandra Orchard, Maria J. Martin, Alex Ignatchenko, Burkhard Rost, Christine A. Orengo, Marc Robinson-Rechavi, Dannie Durand, Steven E. Brenner, Casey S. Greene, Sean D. Mooney, Iddo Friedberg, Predrag Radivojac

## Abstract

The Critical Assessment of Functional Annotation (CAFA) is a long-standing community effort to independently assess computational methods for protein function prediction, to highlight well-performing methodologies, to identify bottlenecks in the field, and to provide a forum for the dissemination of results and exchange of ideas. In its fifth round (CAFA5) of triennial challenges, a partnership with Kaggle Inc. facilitated participation from a large community of data scientists and computational biologists through a competitive prospective challenge on the crowdsourcing platform. In this work, we present an in-depth analysis of the submitted predictions and report improvements in accuracy over all methods from the previous CAFA challenges. We further introduce a new evaluation setting for proteins with pre-existing (incomplete) annotations and identify the need for methods that better leverage existing annotations to predict those that will be discovered later. Finally, we characterize the prospective evaluation framework by examining performance on a strict set of unpublished annotations and across intermediate database releases. Our results indicate that recent developments in the field, such as the availability of protein language models and accurately predicted 3D structures, as well as the growth of experimental annotations through biocuration, have all contributed to performance improvements. ^1^

## 1 Introduction

Critical assessments have emerged as effective mechanisms for tracking scientific progress and driving advances through simultaneous collaboration and competition [11]. The Critical Assessment of protein Structure Prediction (CASP) was the first such effort in molecular biology [23] and has recently led to remarkable developments in 3D structure prediction [34, 35, 8, 21]. The success did not come suddenly or easily [22]. Over two decades of meticulous method development, refinement of assessment metrics, community building, sustained growth of experimental 3D structures, and recent advances in deep learning were all required to reach current performance levels [8].

The next frontier in computational biology, and a field that can similarly benefit from community-driven assessments, is the prediction of protein function at high resolution and across multiple levels of biological organization. Solving this problem demands even broader collaborative effort among diverse communities, including method developers, biocurators, and experimental scientists [29]. Protein function prediction is not a new problem [30, 36, 13], but it presents specific challenges that distinguish it from other classification tasks. A protein often has multiple functions at different levels of abstraction, from molecular activities to roles in complex biological processes, making the prediction task inherently multidimensional. Experimental annotations contain errors [32] and accumulate in a biased manner [25]. The annotations also remain sparse and incomplete, with most proteins lacking comprehensive functional characterization. This makes it difficult to distinguish between truly absent functions and those simply not yet experimentally validated [12]. The problem complexity, data issues, and continually evolving ground truth complicate both model training and evaluation [17, 18], necessitating robust assessment frameworks that can evaluate predictions against newly accumulated experimental annotations rather than relying on static benchmarks.

The Critical Assessment of Functional Annotation (CAFA) has become a unique platform for independently evaluating algorithms in protein function prediction and has emerged as a forum for exchanging ideas and research contributions [27, 19, 38, 28]. Every three years, the CAFA organizers release a large number of proteins whose functions are unknown or partially known. Model developers then use data and methodology of their choice to generate and submit predictions for these targets by the submission deadline set to occur several months after the target release. The submission is then closed, and the organizers collect the predictions. After a waiting period, the organizers collect protein functions accumulated after the submission deadline and evaluate predictor performance only on newly accumulated annotations. This evaluation is *prospective* since it considers only annotations that were unknown via experimental validation at the time prediction algorithms were developed and trained. The CAFA challenges have generally contributed to the development and exposure of new function-prediction methods as well as to new strategies for performance assessment, thus advancing the field [27, 19, 38, 28].

To promote increased engagement with the wider machine learning community, the fifth CAFA challenge (CAFA 5) was held as a competition in partnership with Kaggle, Inc. Kaggle is a platform for data science competitions with 28 million global users [1]. Since its founding in 2010, Kaggle has hosted hundreds of data science competitions, some of which offer monetary prizes for the best-performing models, in addition to site-specific points awarded for participation and strong performance. In addition to hosting competitions, the website provides resources for learning data science skills, a platform for community discussion, and cloud computation resources. Competitions on Kaggle have been shown to be valuable tools for rapid scientific discovery [37] and can also yield insights into machine learning approaches that can be explored by application-area researchers [4, 20]. The established community and framework provided by Kaggle for attracting data scientists to engage with predictive challenges made it a compelling partner for the CAFA 5 challenge.

In this paper, we present an in-depth analysis of the CAFA 5 challenge results and examine the effects of the crowdsourced competition format on participation and prediction quality. Our contributions are fourfold. First, we expand the evaluation framework by introducing a partial-knowledge setting to assess predictions of new annotations added to proteins with pre-existing functional characterization. This setting accounts for nearly 70% of accumulated annotations and will increasingly dominate future evaluations as annotation databases grow and fully unannotated proteins become rarer. We derive information-theoretic evaluation measures for this setting, extending the information accretion framework to handle conditional information gain. Second, we characterize the prospective evaluation framework through complementary analyses: evaluating predictions on annotations from publications released after the submission deadline, a stricter but smaller evaluation set that may reveal publication mining or distributional shifts, and by tracking performance across intermediate UniProt [3] releases, demonstrating that measured performance varies considerably until sufficient annotations accumulate. Third, by partnering with Kaggle, we achieved a 22-fold increase in participating teams compared with CAFA 4 [28], with participants from 78 countries, representing broad scientific and technical backgrounds. Fourth, we report that top-performing methods demonstrate measurable gains over previous CAFA challenges, especially in the no-knowledge and limited-knowledge settings. In the newly introduced partial-knowledge setting, for which developers did not specifically optimize their models, performance is lower. This presents an important new challenge in protein function prediction and an opportunity for methodological innovation.

## 2 Methods

In CAFA, protein function prediction methods are evaluated through a prospective, time-stratified experimental design. Target proteins are released at a specified time point (*t*_*ℓ*_) of challenge launch, predictions are collected by a submission deadline (*t*_0_), and performance is assessed against functional annotations accumulated during a subsequent evaluation period (ending at *t*_*e*_). This framework ensures that ground truth annotations used for evaluation did not exist during model training, thereby reducing data leakage and providing a realistic assessment of predictive performance. We describe below the experimental timeline and evaluation framework, followed by data collection protocols, assessment metrics, baseline methods, and the integration of crowdsourced participation through Kaggle.

### 2.1 Problem Setting

#### Prediction task and timeline

The prediction task in CAFA 5 involved predicting protein function as described by Gene Ontology (GO) terms [6] across three aspects of function: Molecular Function (MF), Biological Process (BP), and Cellular Component (CC). The Gene Ontology is a concept hierarchy structured as a directed acyclic graph (DAG), where nodes (concepts or functional terms) are connected by relational ties, with more specific terms serving as descendants of broader, more general terms. Predictions must be made on the ontology structure frozen at the target release time point *t*_*ℓ*_. We refer to this ontology as 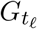.

At time *t*_*ℓ*_, a target set 𝒯 of |𝒯| = 141,865 proteins was released for which participants were invited to predict functional annotations. These were proteins that lacked experimentally validated GO annotations in at least one GO aspect from 90 species of interest previously identified by CAFA [27, 19, 38]. Participants were allowed to use any computational approach and any data available up until *t*_0_ to generate predictions. The required output was a set of confidence scores for GO terms for each target protein. All predictions were submitted by the deadline *t*_0_, after which no modifications were permitted. Following submission, annotations accumulated through published experimental research and were added by biocurators to the UniProt Knowledgebase [3]. At evaluation time *t*_*e*_, performance was assessed against experimentally validated annotations added to UniProt between *t*_0_ and *t*_*e*_. Figure 1 illustrates the experiment timeline.

**Figure 1.**
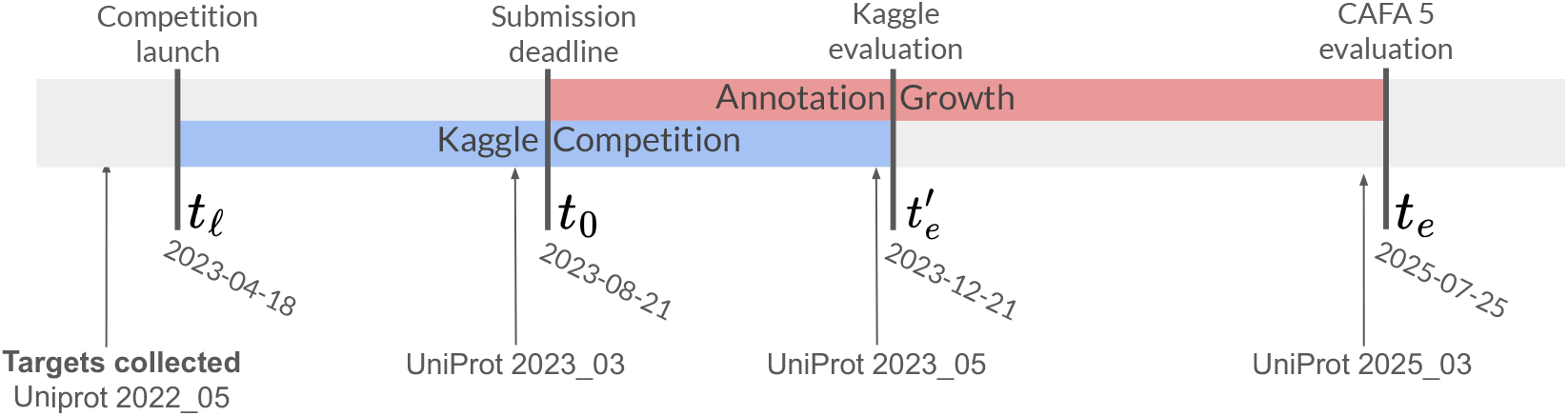
CAFA 5 challenge timeline

#### Evaluation settings

Evaluation was divided into three distinct settings that reflect different prior knowledge for each protein and GO aspect (Figure 2a). The No-Knowledge (NK) setting applies to proteins with no experimentally validated GO annotations in any aspect before *t*_0_. For protein *i* ∈ 𝒯, we evaluate predictions across any aspects if new annotations are added during [*t*_0_, *t*_*e*_]. Limited-Knowledge (LK) evaluation captures the performance on proteins with experimentally validated annotations in at least one GO aspect before *t*_0_, but lacking annotations in one or more aspects. For protein *i* ∈ 𝒯 with prior annotations in aspect(s) *A* but not in aspect(s) *B*, we evaluate predictions only in the previously unannotated aspects *B* if new annotations are added to those aspects during [*t*_0_, *t*_*e*_]. Partial-Knowledge (PK) setting evaluates predictions on proteins with existing experimentally validated annotations in an aspect before *t*_0_ that receive additionalannotations in the same aspect during [*t*_0_, *t*_*e*_]. For protein *i* ∈ 𝒯 with prior annotation set 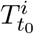 inaspect *A*, we evaluate predictions only on the newly added terms 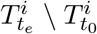. A protein can be included in both the LK and PK evaluations for different GO aspects. The PK setting was introduced in CAFA 5 to address the growing proportion of proteins with partial functional characterization. As annotation coverage increases across iterations, fewer proteins remain completely unannotated, making PK evaluation increasingly important for assessing a method’s ability to refine and expand existing functional knowledge (Figure 2b).

**Figure 2:**
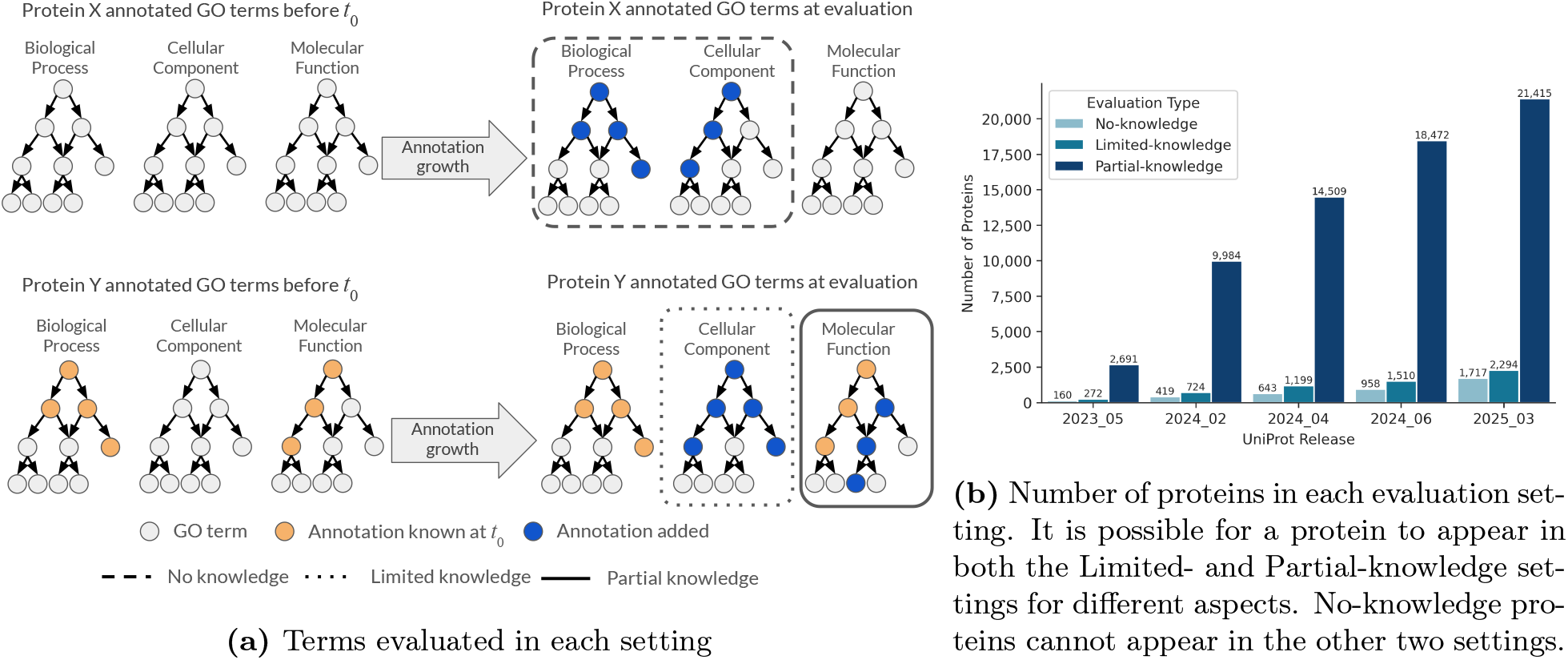
Evaluation settings. (a) Annotations for a protein across three GO aspects. If a protein had no annotations before *t*_0_ (top left) and has annotations added during the annotation growth period (top right), it will be included in the No-Knowledge (NK) evaluation. For proteins with annotations before *t*_0_ (bottom row), aspects with no previous annotations will be included in the Limited-Knowledge (LK) evaluation (aspect inside dotted line). Aspects that had annotations before *t*_0_ and new annotations added will be used in the Partial-Knowledge (PK) evaluation (aspect inside solid line)). Only new annotations (dark blue nodes) will be included in the evaluations. (b) As annotations grow, more Partial-Knowledge terms accumulate in the target set of proteins.

#### Metrics

Predictions were evaluated using several key metrics. By treating the prediction task as a collection of binary predictions, precision (*pr*), recall (*rc*), and *F*_1_-score were computed. The information-theoretic counterparts of these measures, weighted by information accretion (IA) [10], were used to incorporate the hierarchical relationships between GO terms and the amount of surprise of observing each term. We report both micro-averaged (*µ*) metrics (pooling all predictions across proteins) and macro-averaged (*M*) metrics (averaging per-protein performance).

For a protein *i* ∈ 𝒯, let *P*^*i*^(*τ*) be the set of terms in one aspect *A* of the GO DAG (*A* ⊂ *G*) for which the prediction algorithm yields a score greater than or equal to *τ* ∈ [0, 1]. Let 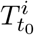 and 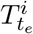 be the set of terms for which there are experimentally validated annotations at *t*_0_ and *t*_*e*_, respectively. Let 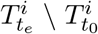 be the *new* annotations added to the aspect *A* for protein *i* between time *t*_0_ and *t*_*e*_; *i*.*e*., the ground truth annotations on which evaluation will be calculated. Sinceonly annotations on these newly added terms will be evaluated, only predictions in 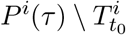 are considered. For notational compactness, we write simply *P*^*i*^(*τ*) for the predicted annotations and 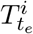 for ground truth annotations. For NK and LK evaluation, *T*_*t*_ = ∅ and thus 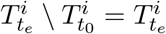 and 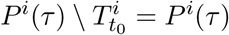. We compute macro- and micro-averaged precision *pr* and recall *rc* as follows:

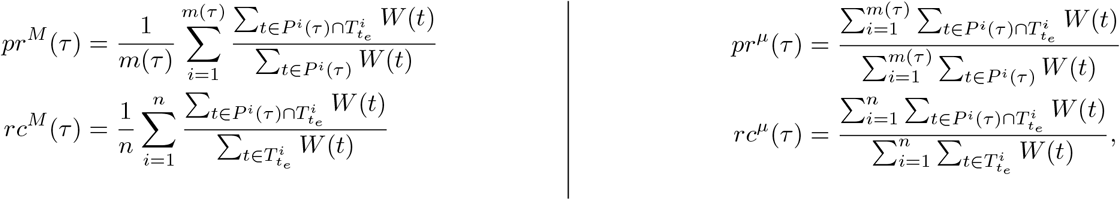

where *m*(*τ*) is the number of proteins with at least one term *t* with model score greater than or equal to *τ* and *n* is the number of proteins with at least one term in the ground truth annotations. We define the coverage (*cov*), of a predictor as the proportion *m*(*τ*)*/n*. The *F*_max_ score is defined as *F*_max_ = max_*τ*_ *F*_1_(*τ*), where 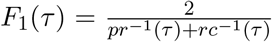at threshold *τ*, providing a single-number summary of prediction quality at the optimal threshold. The weighting function *W* (*t*) is the indicator function 𝟙(·) for the standard precision and recall computation and is the information accretion IA(*t*) for weighted metrics.

IA represents the information content of term *t*, computed and frozen at time *t*_0_ to ensure consistent weighting throughout the evaluation. The information accretion of a set of terms 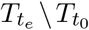 is defined as

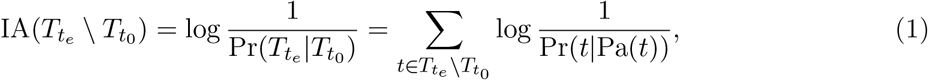

where Pa(*t*) denotes the parent terms of *t* in the ontology. The information accretion weights terms by their rarity in the annotated protein corpus, down-weighting the importance of common, less informative terms. For NK and LK evaluations where 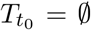, this reduces to the unconditional information content IA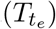 [10]. For the PK setting, IA accounts for the conditional information of newly added terms given the prior annotations 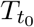; see Appendix A for derivation.

We also compute the semantic distance metric 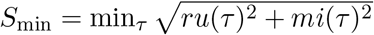, which measures the remaining uncertainty (*ru*) and misinformation (*mi*) in predictions relative to ground truth, providing complementary information to precision-recall based metrics [10].

### 2.2 Crowdsource Challenge

CAFA 5 partnered with Kaggle to host the challenge as a public competition. In addition to continuing to engage the protein function prediction community, this format facilitated the introduction of the problem to data scientists and machine learning practitioners who may not traditionally engage with computational biology research. Bringing these communities together to address this prediction challenge promotes diversifying the approaches applied to the problem.

#### Design adaptations

Hosting CAFA 5 on Kaggle required several adaptations to the traditional CAFA format to align with the platform’s structure and user expectations.

#### Training Dataset

We provided participants with a curated training dataset consisting of all 142,246 proteins with GO annotations validated by experimental evidence codes (Appendix B.2) in the UniProt Knowledgebase [3] version 2022_05.

#### Simplified Evaluation Metric

While CAFA evaluations traditionally employ multiple metrics, including standard and weighted precision, recall, *F*_max_, and semantic distance, to promote comprehensive insights, the Kaggle competition required a single metric for ranking participants. We selected the IA-weighted *F*_max_ score, averaged across the three GO aspects and included only NK and LK annotations.

#### Intermediate Ground Truth

To provide ongoing feedback during the four-month competition period (April–August 2023), we established a set of intermediate ground truth annotations. Kaggle uses a leaderboard that gives participants feedback on their relative performance, thereby stimulating further engagement. To allow for this option, we have worked with UniProt to secure a set of curated proteins that were not publicly available in the database at *t*_*ℓ*_ and were held out until the close of the competition at *t*_0_. This intermediate ground truth contained 76 proteins with 2,382 new annotations.

#### Preliminary evaluation

To award prizes for the competition, an initial evaluation was conducted at 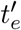, December 2023 (Figure 1). While this allows for the accumulation of some new annotations, this condensed timeline provides fewer ground truth annotations than the full evaluation. The preliminary evaluation was conducted on 10,304 annotations accumulated for 499 proteins in the target set between 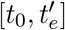.

#### Participation

The Kaggle format resulted in unprecedented participation levels. During the four-month challenge, 1,625 teams from 78 countries submitted at least one prediction, resulting in a total of 2,850 models (Figure 3). This represents a 22.2-fold increase in participating teams compared to CAFA 4 and a 17.6-fold increase in submitted models. Geographic diversity increased substantially. Comparatively, CAFA 3 had co-authors from institutions in 21 countries, and none of which were from South America, sub-Saharan Africa, or Oceania, all of which were represented in CAFA5 teams (Figure 3b). The record participation demonstrates the interest in function prediction to a wider machine learning audience and the potential for engagement with that community to make computational advancements in this field. After the competition, top-performing teams were asked to present their approaches, providing insights and ideas to be publicly shared [2].

**Figure 3.**
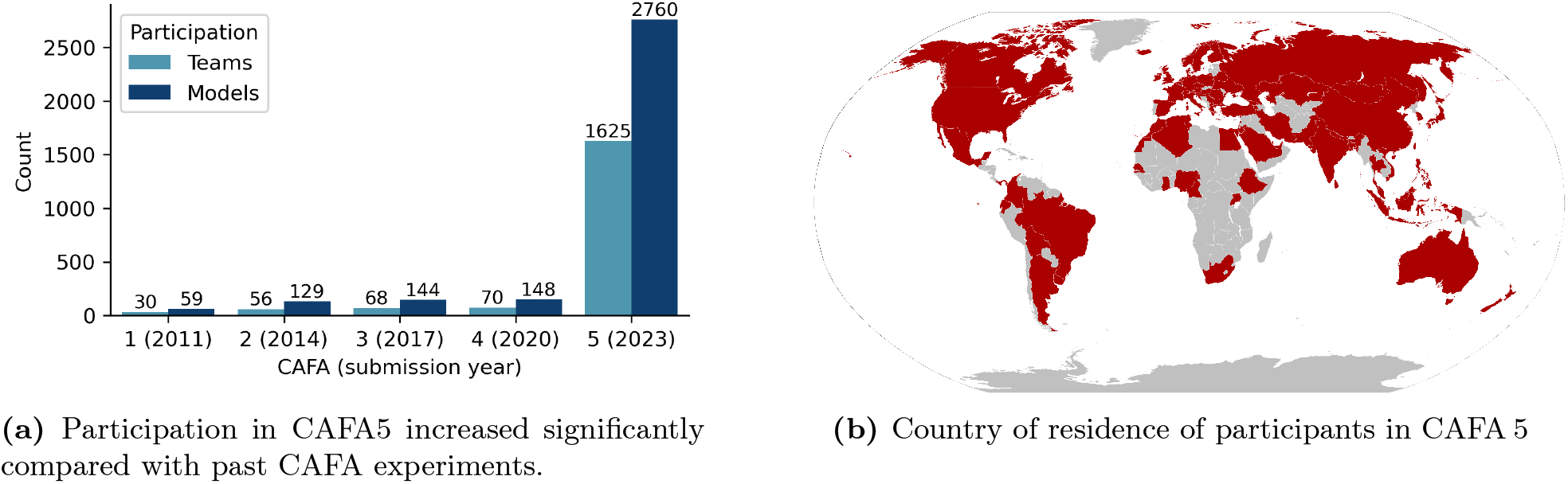
(a) Participation increased significantly compared to past CAFA experiments. (b) Participants from 78 countries registered and submitted predictions in CAFA 5.

### 2.3 Evaluation Data

The final CAFA 5 evaluation was conducted using annotations from the UniProt Knowledgebase release 2025_03, collected 23 months after the submission deadline *t*_0_. All predictions were made on the GO structure frozen at *t*_*ℓ*_ (version 2023-01-01), and known annotations 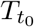 at *t*_0_ were filtered using UniProt release 2023_03. We considered only experimentally validated annotations (evidence codes listed in Appendix B.2) for both known annotations at *t*_0_ and the evaluation ground truth at *t*_*e*_. All annotations were propagated according to the GO rules of the relations *is a, part of, regulates, negatively regulates*, and *positively regulates*. That is, if a protein is annotated with term *t*, it is implicitly annotated with all ancestors of *t* in the GO DAG.

From the original target set 𝒯, proteins were excluded from the final evaluation if they were removed from UniProt of if their amino acid sequence changed. Specifically, 42 and 18 proteins were excluded because they were removed from UniProt or merged with other entries, respectively, between *t*_*ℓ*_ and *t*_*e*_. Additionally, 598 proteins were removed because their amino acid sequences in UniProt changed. Because sequence-based methods may produce different predictions for altered sequences, we excluded these proteins to maintain consistency between training and evaluation conditions. After filtering, the final evaluation set consisted of 24,405 proteins and 234,529 annotations. Table 1 summarizes the composition of the evaluation dataset, stratified by GO aspect (MF, BP, CC) and evaluation setting (NK, LK, PK). As shown in Figure 2b and in Table 1, the PK setting dominates the evaluation, accounting for 69.7% of accumulated annotations.

**Table 1.** Counts of proteins and terms used for evaluation.

| Evaluation type | Proteins | Terms |  |  |  |
| --- | --- | --- | --- | --- | --- |
|  |  | BP | CC | MF | Total |
| No Knowledge | 1,717 | 21,033 | 10,343 | 4,182 | 35,558 |
| Limited Knowledge | 2,294 | 23,046 | 7,943 | 4,633 | 35,622 |
| Partial Knowledge | 21,415 | 117,331 | 54,528 | 23,963 | 163,349 |
| Total | 24,405 | 161,410 | 42,249 | 30,870 | 234,529 |

We also construct datasets following the same protocols on UniProt release versions 2023_05, 2024_02, 2024_04, and 2024_06. We use these datasets to show the change in performance as the annotations grew. Details on these data can be found in Table A2.

### 2.4 Annotations from New Publications

The prospective evaluation in CAFA relies on new, previously unknown annotations being added to the UniProt database as those annotations are discovered or confirmed experimentally and published in the literature. However, the date of addition to UniProt is not always a reliable proxy for date of first discovery or experimental validation. We therefore analyzed the publication date of source publications for all annotations added to UniProt since *t*_0_ (2023-08-21) which had corresponding PubMed or DOI identifiers. To construct a stringent prospective evaluation of annotations unknown in the literature before the annotation growth phase, we filtered out all annotations with no reference or publication associated in UniProt and any published before before *t*_0_ (2023-08-21). We found that only 3,627 annotations out of 68,940 unique annotations (5.3%) added to UniProt since *t*_0_ associated with CAFA target proteins had publications corresponding to dates *after t*_0_.

We conducted a complementary evaluation of CAFA5 methods for only the subset of annotations attributed to publications after *t*_0_. This set of 1,568 proteins (23,317 propagated annotations) was used for a prospective evaluation where the annotations did not exist in UniProt during model development *nor did the publications on which the annotation is based*. Table 2 describes the dataset used in this evaluation, showing that these annotations make up 9.9% of the full evaluation dataset.

**Table 2.** Dataset counts for head-to-head (CAFA 2-5) evaluation and unpublished annotations evaluation.

| Evaluation type | Dataset: Head-to-head |  |  |  |  | Dataset: Unpublished |  |  |  |  |
| --- | --- | --- | --- | --- | --- | --- | --- | --- | --- | --- |
|  | Proteins | Terms |  |  |  | Proteins | Terms |  |  |  |
|  |  | BP | CC | MF | Total |  | BP | CC | MF | Total |
| No knowledge | 1,102 | 13,399 | 5,771 | 2,072 | 21,242 | 272 | 3,540 | 1,222 | 562 | 5,324 |
| Limited knowledge | 1,769 | 18,586 | 4,819 | 2,850 | 26,255 | 351 | 4,016 | 726 | 1,074 | 5,816 |
| Partial knowledge | 15,732 | 87,100 | 14,921 | 15,126 | 117,147 | 1,073 | 8,689 | 2,298 | 1,190 | 12,177 |
| Total | 17,818 | 119,085 | 25,511 | 20,048 | 164,644 | 1,568 | 16,245 | 4,246 | 2,826 | 23,317 |

### 2.5 Comparing Against Past CAFA Challenges

To assess progress of the field over time, we conducted a head-to-head evaluation of CAFA5 methods compared to previous CAFA challenges [19, 38, 28]. We excluded CAFA 1 due to the limited overlap between the GO used at that time and the current ontology. To construct an evaluation set, we found the set of proteins that appear in the target sets of all CAFA challenges, 𝒯^CAFA 5^∩ 𝒯^CAFA 4^ ∩ 𝒯^CAFA 3^ ∩ 𝒯^CAFA 2^, where the superscript indicates the CAFA version. We evaluated predictions only on common terms across all four GO graph structures.

### 2.6 Baselines

To benchmark performance of predictors, we present four baseline predictors, including Naive, non-experimentally validated evidence, BLAST similarity, and embedding-based similarity.

#### Naïve

The Naive [9] prediction baseline assigns each term in the ontology a score equal to its frequency in the database. To generate these scores, we filtered the annotations available at *t*_0_ (GOA 2023-07-12) annotations to keep only those that were experimentally validated. We propagated these terms on the ontology available at *t*_0_ (GO 2023-07-27) to get the full set of annotations. We kept only the terms in the target ontology (GO 2023-01-01) and finally propagated the remaining terms on the target ontology to ensure connectivity in the graph. The scores correspond to the number of proteins with each of the terms annotated divided by the number of proteins in the set.

#### Non-Experimentally Validated Evidence

We predicted terms with a score 1 if the term was annotated at *t*_0_, regardless of evidence code. This baseline shows how well non-experimentally annotations predict future experimentally validated annotations. The performance of this baseline shows how accurately UniProt electronic, computational, and phylogenetically inferred evidence codes predict experimentally validated functions.

#### BLAST

The BLAST baseline method employed NCBI BLAST+ version 2.15.0 to search target sequences against a reference database of proteins with experimentally verified GO annotations available at *t*_0_ [5]. This database comprised 137,372 proteins retrieved at *t*_0_ (GOA 2023-07-12), with sequences obtained from UniProt release 2023_03. Of these, 84,562 proteins originated from the Swiss-Prot subset and 52,810 from TrEMBL. BLASTp searches were performed using default parameters, and functional annotations were transferred from reference database hits to target proteins based on sequence similarity. For each GO term, the confidence score was determined by the maximum local sequence identity observed between the target protein and any experimentally annotated reference protein bearing that term. When multiple reference proteins were annotated with the same GO term, the highest sequence identity score was retained.

#### Embedding Similarity

Similarly to the BLAST baselines, we were interested in transferring annotations from *t*_0_ based on sequence similarity for each target sequence, from the same reference database used for the BLAST baseline. Unlike BLAST, for the protein language model (pLM) embedding baseline, we used the similarity between learned embedding representations. The embeddings are vector representations used in a pLM model internally to learn the language of amino acid sequences, based on observed training sequences. We used ProtT5 embeddings [15] from the prot_t5_xl_half_uniref50-enc model downloaded from Hugging Face. For the similarity measure, we used cosine and Euclidean distances between per-protein embeddings which has been shown to capture relevant protein relationships [7, 16, 33, 24]. Two normalization approaches were implemented depending on the similarity metric. Euclidean distances were converted to similarity scores through normalization: similarities were calculated as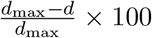, where *d* represents the Euclidean distance between a target and reference set protein, and *d*_max_ is the maximum distance observed across all targets. For cosine similarity, min-max scaling was applied to transform the similarity range to [0, 100]. Both normalization methods ensure comparable similarity scores regardless of the underlying distance metric. We constructed baselines using three- and five- nearest neighbors, based on the embedding distance, and used the best-performing as the reported baseline.

#### Head-to-Head BLAST

For comparing to past CAFA experiments, we used a separate BLAST predictors where the database of proteins from which query (target) proteins were combined come from the annotations available before the close of the corresponding CAFA: 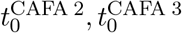, and 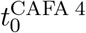 for CAFA 2, 3, and 4, respectively; see Table A2. BLAST annotations from the databases of increasing size can be used as a proxy to understanding how much of the improvement in the prediction comes from growing annotation data versus the improvements in modeling.

## 3 Results and Discussion

### 3.1 CAFA 5 Models Improve Over Previous Models

We evaluate the performance of top CAFA 5 methods compared to those submitted to previous CAFA iterations to evaluate the progress of the field and the robustness of previous methods. We evaluate each of the top models for CAFAs 2-5 on the portion of the CAFA 5 evaluation data that overlaps with previous targets, as described in Section 2.5. This evaluation is also the first to evaluate previous CAFA models on the new partial-knowledge evaluation setting. In all cases, we consider only the best-performing model from each team. Proteins in the evaluation set were sampled with replacement and performance metrics were computed for 5,000 bootstrap samples [14]. We find that top methods in CAFA 5 outperform all previous methods with respect to 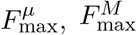, and *S*_min_ for all evaluation settings and GO aspects with only exception being the BPO aspect under partial knowledge (PK) evaluation with respect to 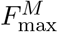(see Figures 4 and A2).

**Figure 4.**
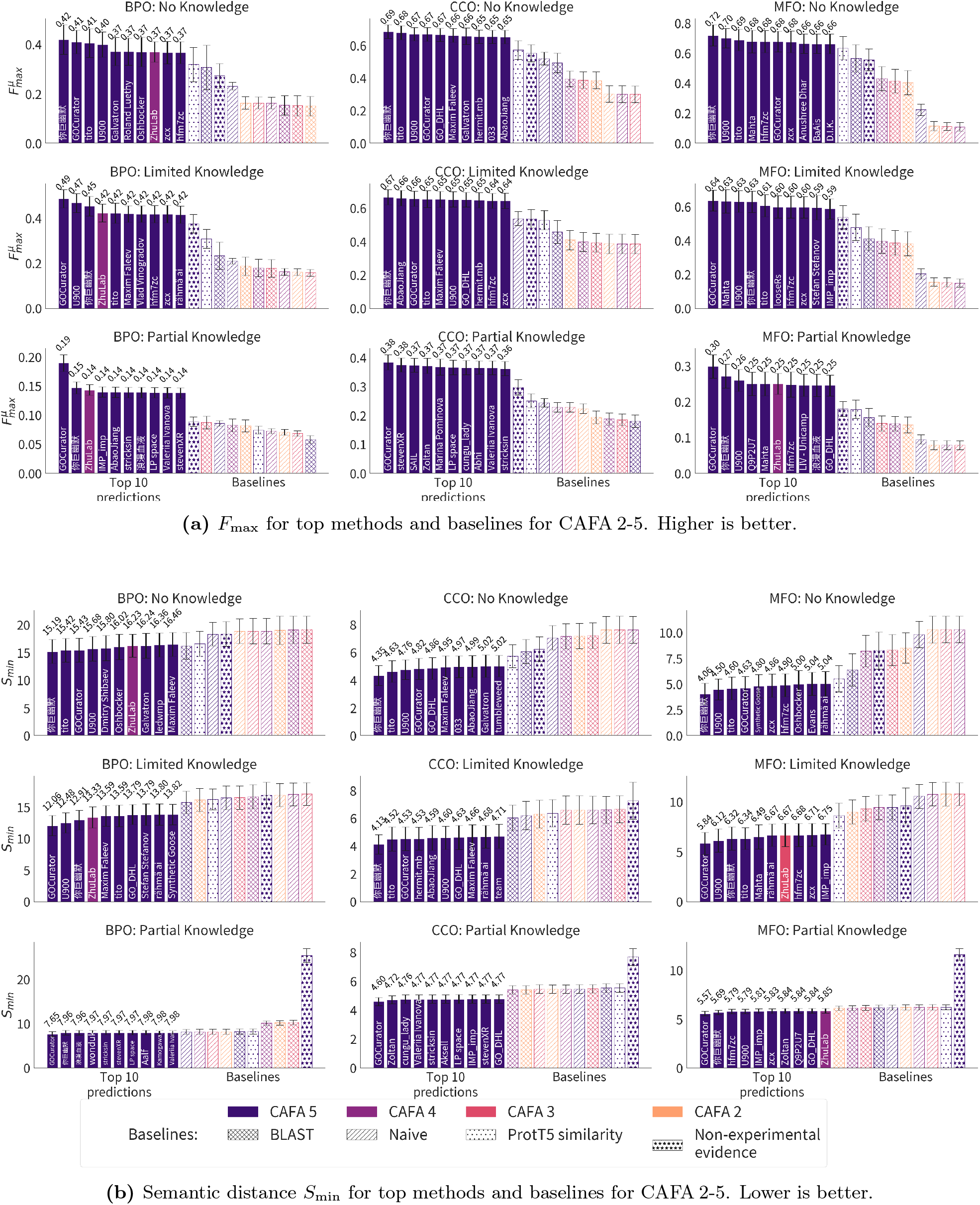
Head-to-head comparison: Error bars show 95% confidence intervals for 5,000 bootstrap samples. Group names are shown inside bars and bar colors indicate CAFA experiment (2-5). CAFA 5 methods outperform previous top methods in all aspects and evaluation settings. Only one team from previous CAFA experiments (3 and 4) perform in the top 10.

There are a few models from previous CAFA challenges that reach top-ten performance on the CAFA 5 evaluation data. The top-performing team from CAFA 4 [28], ZhuLab, performs among the top ten methods with respect to 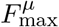 for all settings for the BPO and for MFO aspects for the PK setting. This team also reaches top-ten performance with respect to 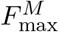 in the BPO and MFO PK evaluations (Figure A2) and with respect to the *S*_min_ metric for BPO in the no-knowledge (NK) and limited knowledge (LK) settings and MFO PK setting. The same team reaches the top ten with their CAFA 3 model [38] for MFO LK with respect to *S*_min_ and BPO PK with respect to 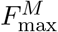. No other teams placed among the top models when compared with CAFA 5 except for the BPO PK evaluation with respect to 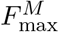. Here, several CAFA 4 models and one CAFA 3 model reach the top ten, shown in Figure A2.

In all cases, the top-ten methods outperformed all baseline methods. Generally, the BLAST and Naive baselines using CAFA 5 look-ups outperformed their earlier counterparts. For 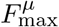, the non-experimental annotations in UniProt were among the best performing baselines. This indicates that non-experimental annotations are fairly reliable, though predictive models from CAFA submissions are able to improve on this baseline. The non-experimental annotations result in higher *S*_min_ measures (worse performance), especially in the PK setting. This reflects mostly a high measure of *misinformation*. This performance drop may indicate that annotations that have non-experimental evidence codes have not *yet* been experimentally validated, but may be in the future. Reevaluation of this baseline will serve to assess the snapshot of non-experimentally validated annotations at the time of CAFA 5 (*t*_0_) as more annotations are tested experimentally.

### 3.2 Partial Knowledge Performance Lags Behind Other Settings

We evaluate all 2,760 models submitted for the CAFA 5 challenge on the full set of experimentally validated annotations added to UniProt since *t*_0_ (2023-08-21). We calculate the performance with respect to 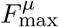, *S*_min_, and 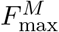. We estimate 95% confidence intervals using bootstrapping on 5,000 iterations [14] and show average results in Figures 6, 8, and A4. Results for the complete precision-recall (PR) curves are shown in Figures 5, 7, and A3. The top ten methods and baseline methods are highlighted in the PR curve plots, but lines for the top 100 methods are also included. Top 100 methods generally outperform baseline methods and in most cases (except CCO), the top few methods have a clear performance improvement over all other methods.

**Figure 5.**
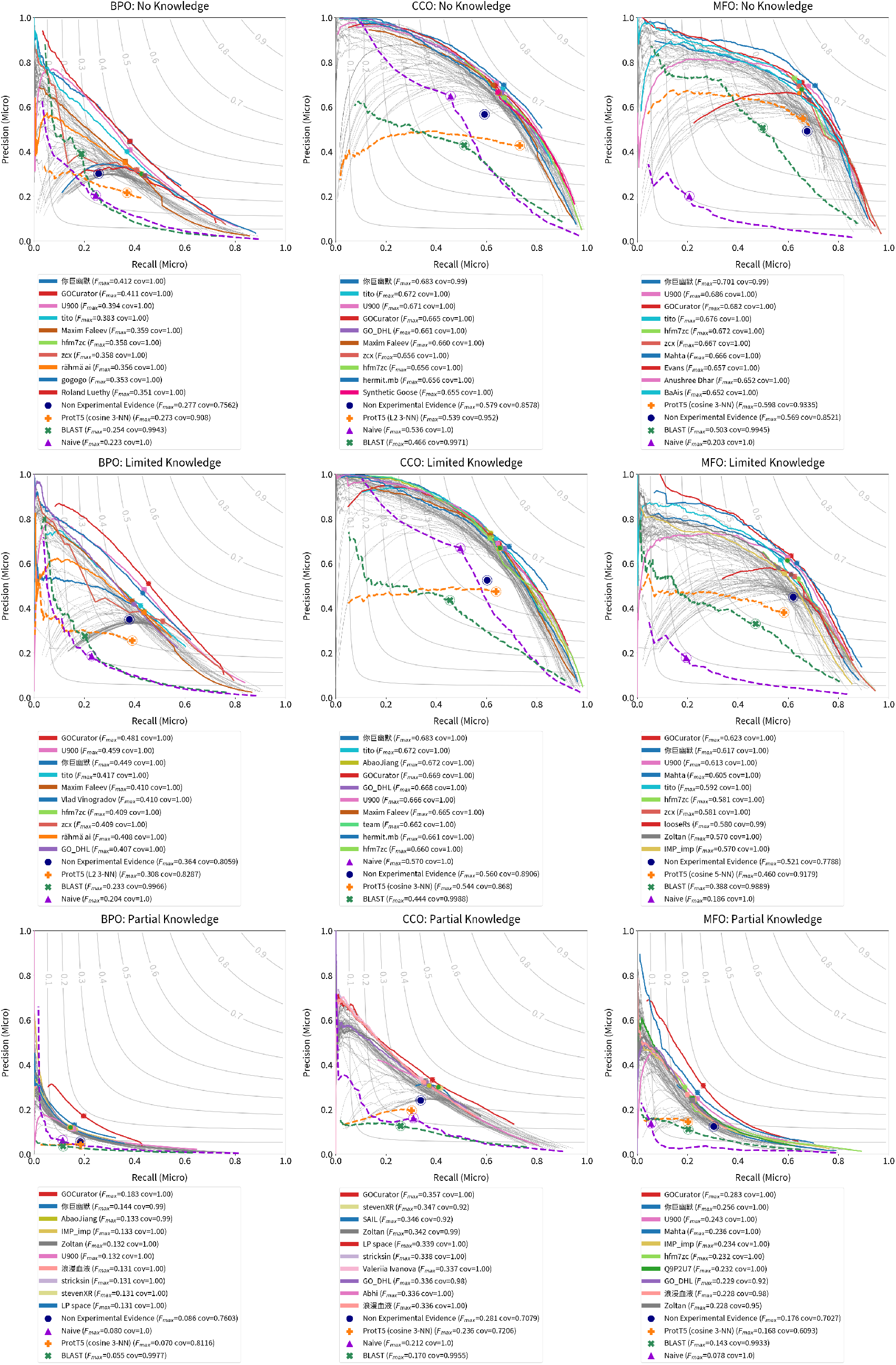
Micro-average precision/recall curves for top 100 submitted methods and baseline methods. The top 10 submitted methods are highlighted in colors and listed in the legend in order of *F*_max_. Higher values, towards upper-right, are better.

**Figure 6.**
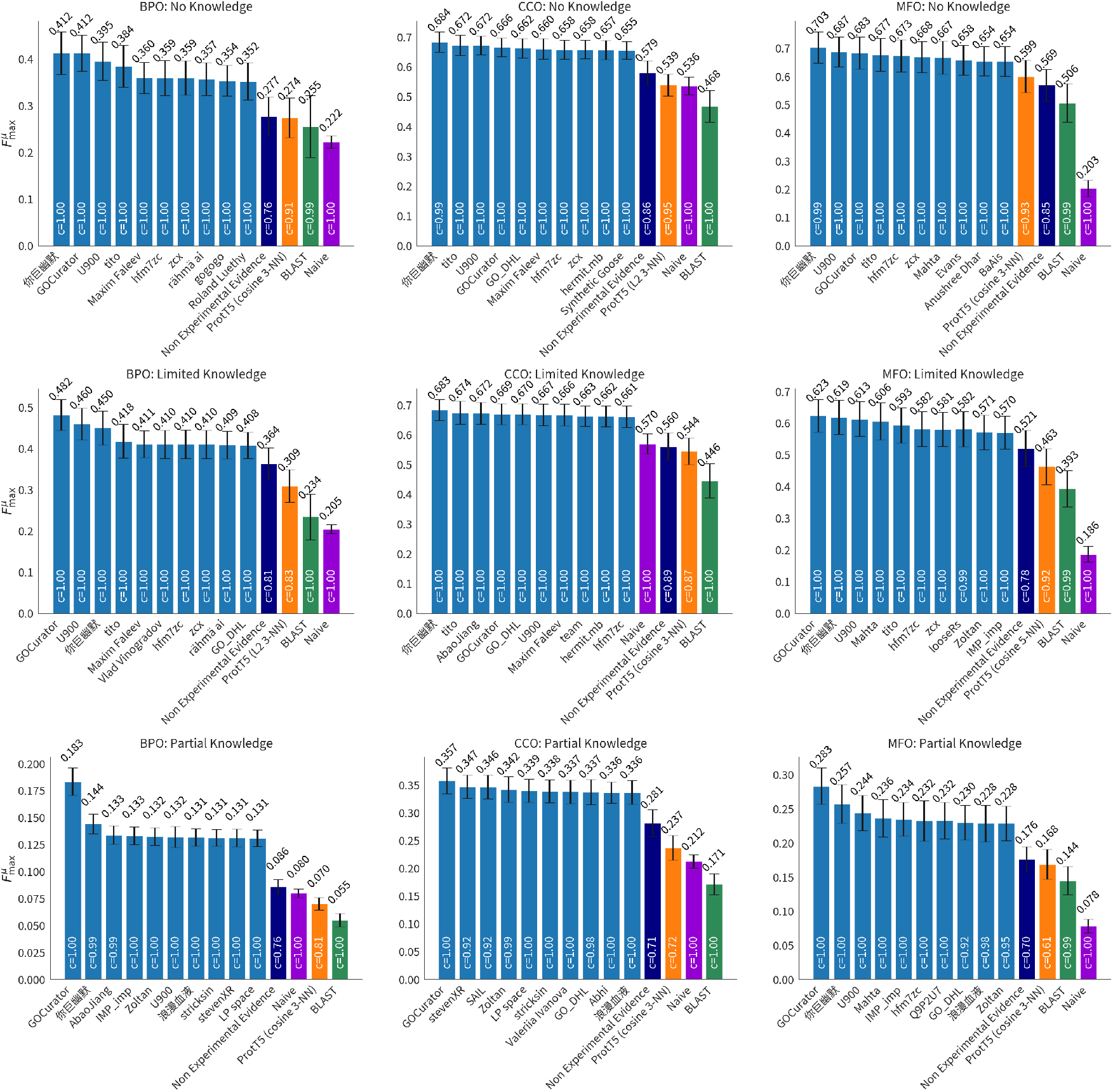
Micro-average *F*_max_ over 5,000 bootstrap samples with 95% confidence intervals. The mean value is shown above the bars. Higher values are better.

**Figure 7.**
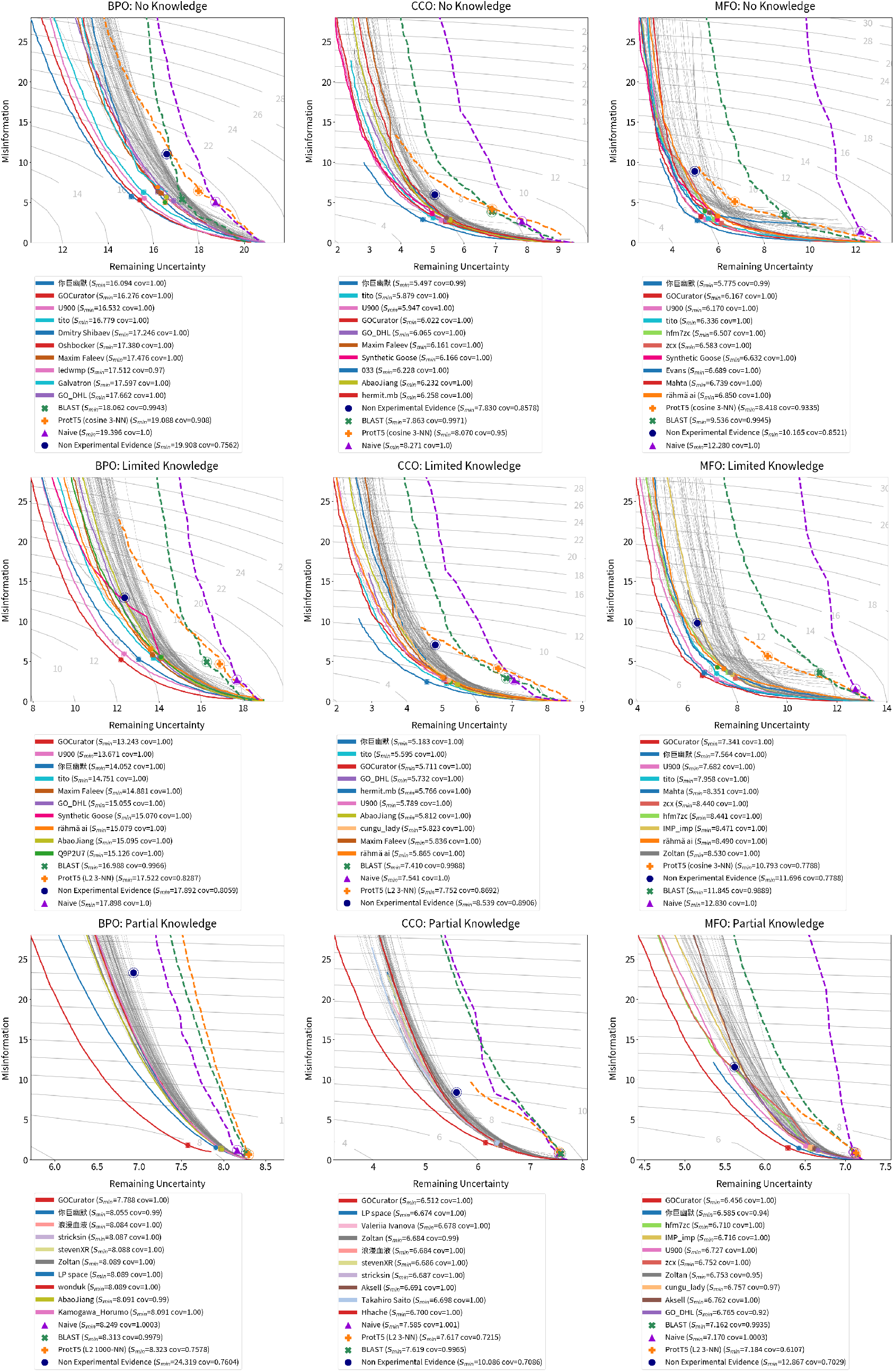
Misinformation/remaining uncertainty curves for top 100 submitted methods and baseline methods. The top 10 submitted methods are highlighted in colors and listed in the legend in order of *S* −score. Lower values, towards lower-left, are better.

**Figure 8.**
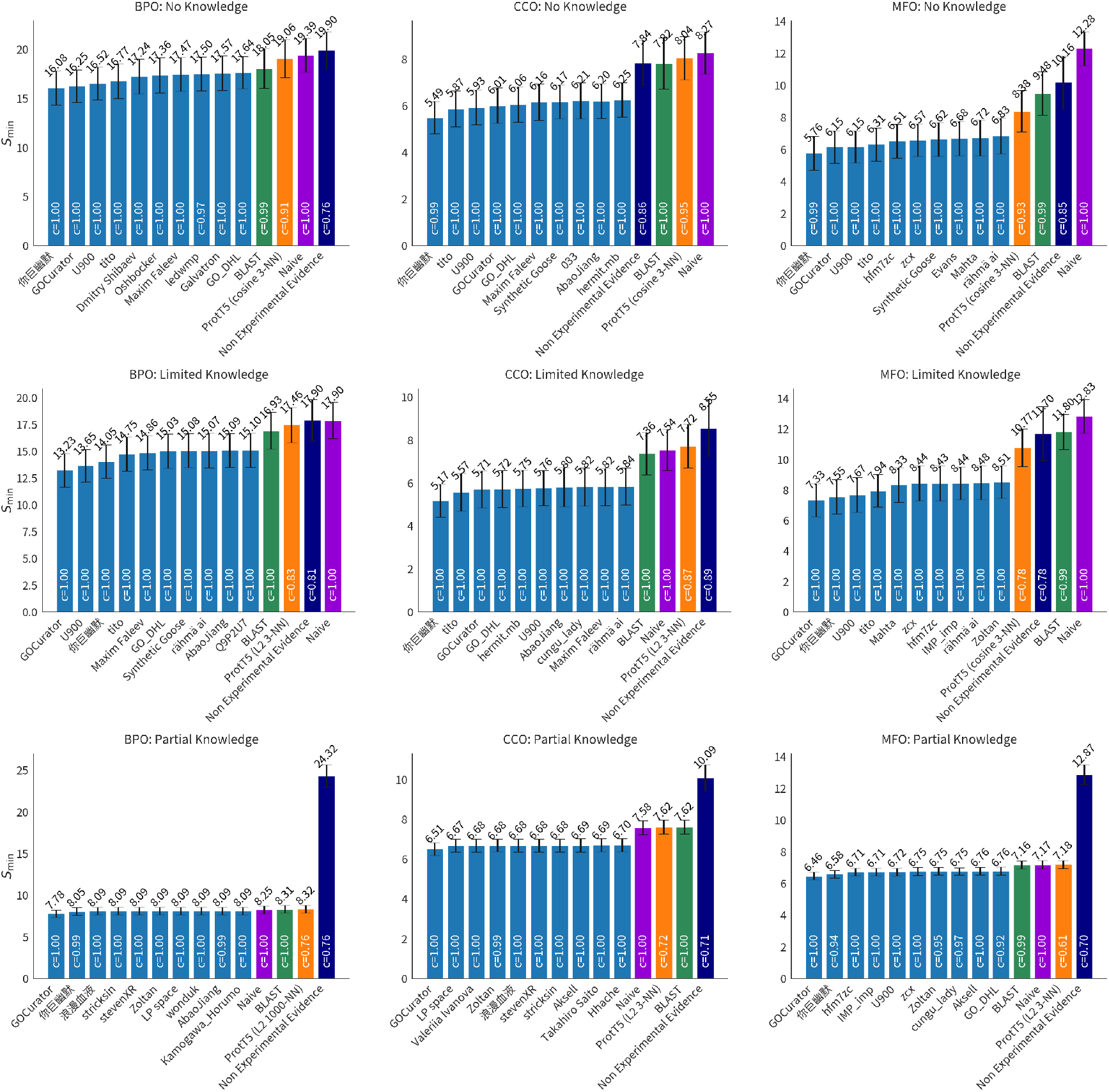
*S*_min_ over 5,000 bootstrap samples with 95% confidence intervals. The mean value is shown above the bars. Lower values are better.

Performance for each GO aspect (BPO, CCO, MFO) is similar for no-knowledge (NK) and limited-knowledge (LK) evaluation for all three metrics, 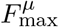, *S*_min_, and 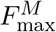. For all three metrics, the performance on the partial-knowledge (PK) evaluation setting is worse. Comparing the NK and PK best-performing model for each aspect, BPO, CCO, and MFO, we see a 55.6% 47.8% 59.7% decrease in 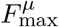, respectively. Comparing LK and PK, we see a 62.0%, 47.7%, and 54.6% decreasein 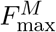. Similar drop in performance is seen in the *S*_min_ increase for models, except for MFO where the performance drop is less severe, and in the case of MFO LK versus PK evaluation, *S*_min_ decreases (improves) by 11.9% for the top model.

The general drop in performance for the PK setting may reflect the increased challenge in the prediction task for terms deeper in the ontology. The difficulty may be attributed to these functions being more difficult to predict or due to the distributional difference between terms added to proteins with prior known terms compared to those added for proteins with no known functions. However, it is not possible to disentangle these potential reasons from model developers not optimizing for this new setting. Since this is a newly introduced setting for evaluation, it may be that models purposely or inadvertently perform worse for such annotations in order to optimize for the prediction task historically evaluated in CAFA challenges. Still, the disparity in performance suggests new directions for improving models by either conditioning on already known annotations, or by focusing on this setting which accounts for the majority of new annotations in UniProt (69.6%).

### 3.3 Evaluation on Unpublished Annotations

The prospective evaluation framework used by CAFA attempts to prevent data leakage between model training and the evaluation datasets. However, UniProt annotations are often gathered from publications and then added to the database. In order to further limit data leakage, we also perform a reevaluation of CAFA 5 models on the subset of annotations that were unpublished at *t*_0_ (2023-08-21). We show the comparative performance for the full evaluation versus the unpublished annotations for 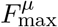 in Figure 9 and for *S*_min_ in Figure A5.

**Figure 9.**
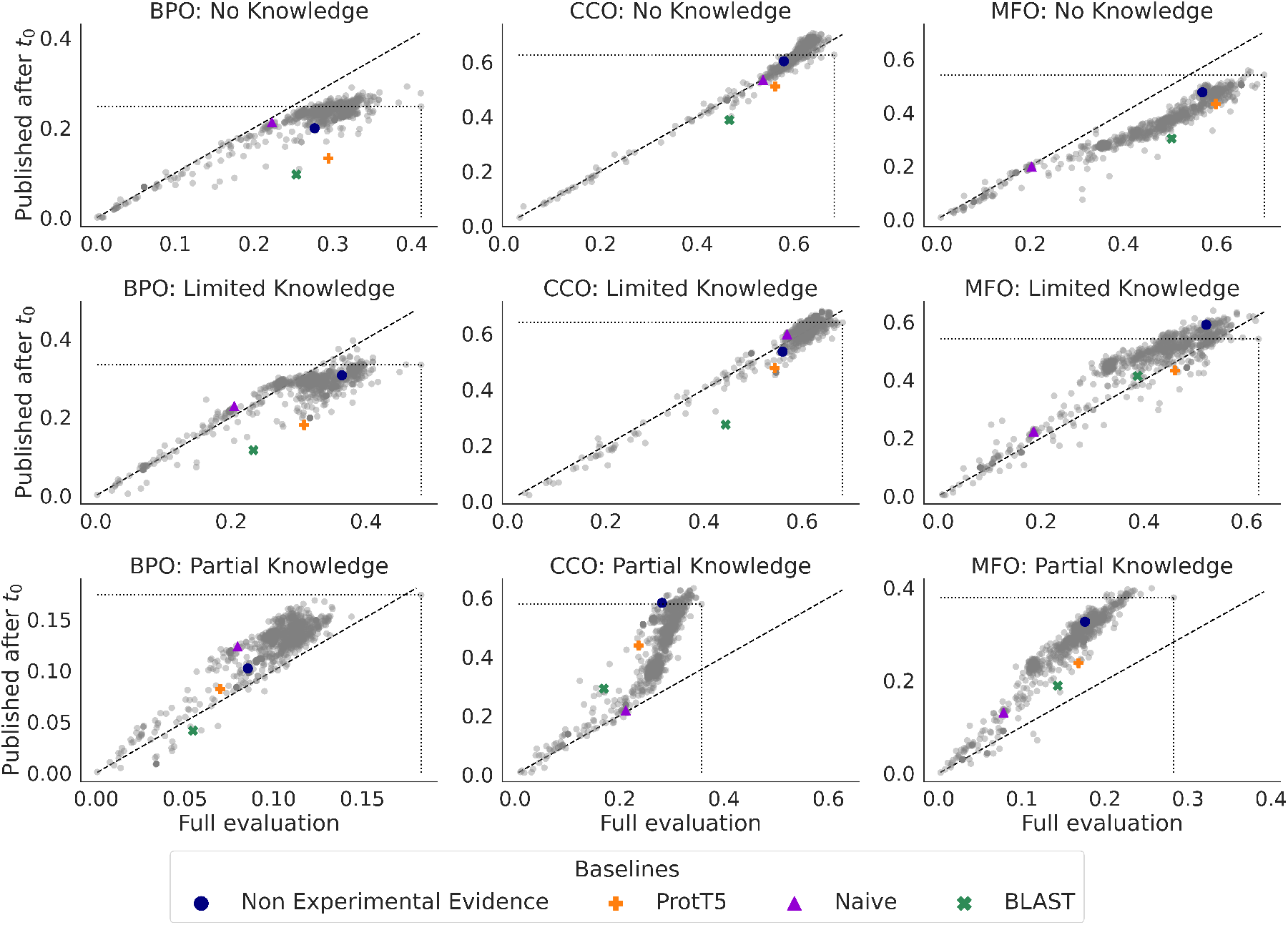
We compare 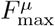 (micro-average) of methods on the full evaluation set (horizontal axis) with a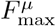 on just the subset of annotations from publications released after *t*_0_ (vertical axis). Points above the diagonal line indicate that the method’s 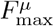 improved when evaluated on the subset of annotations from new publication. Points below the diagonal line indicate a reduction in 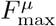. The 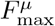 of the topperforming method in the full evaluation (furthest to the right) is designated by a dotted line. In each case except BPO partial knowledge, the top-performing model on the full evaluation is not the best-performing when evaluated on newly-published annotations, as indicated by points lying *above* the horizontal dotted line.

For 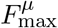, results for the CCO aspect in both no-knowledge (NK) and limited-knowledge (LK) settings remain fairly unchanged in both cases, with most models performing the same and points lying along the diagonal line of Figure 9. Conversely, for the BPO aspect in the NK and LK settings, models tend to perform worse for the set of annotations from publications after *t*_0_. The MFO aspect shows a mix, with the NK setting showing a decline in performance and the LK setting showing a slight improvement. In all aspects, the partial-knowledge (PK) setting shows that models perform better on the newly-published annotations than the full set of annotations.

The evaluation on this set may be a better proxy for performance of GO term prediction on terms that were actually unknown before *t*_0_ since they did not exist in UniProt nor published. However, it is important to note that the set of unpublished annotations is much smaller, about 10% the size of the full evaluation. Therefore, it may not be representative of the distribution of GO terms. It remains to be seen if the patterns of performance on this set will hold as more annotations accumulate. Repeated evaluation with a larger set of unpublished annotations in future CAFA challenges is needed to test the long-term performance of these models.

### 3.4 Effects of Evaluation Over Time

Lastly, we validate the prospective evaluation by repeatedly evaluating the CAFA 5 models on successive UniProt releases between *t*_0_ and the final evaluation time, *t*_*e*_. Figure 10 shows the performance of the top 100 models and baselines with respect to 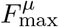. In general, the performance of models evaluated at the first UniProt release after *t*_0_ (release 2023_05) is higher than the performance evaluated in subsequent releases. The 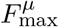 measure drops the most at the second evaluation (release 2024_02) and then changes less for the remaining releases. This pattern may be due to the fact that, as more annotations accumulate, they reflect a more representative distribution of functions, matching more closely the data on which the models were trained. Early on, there are fewer annotations and the performance fluctuates more. As more annotations accumulate, the measure is more robustly evaluating the performance. Similar patterns are shown for *S*_min_ and 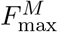(see Figure A6).

**Figure 10.**
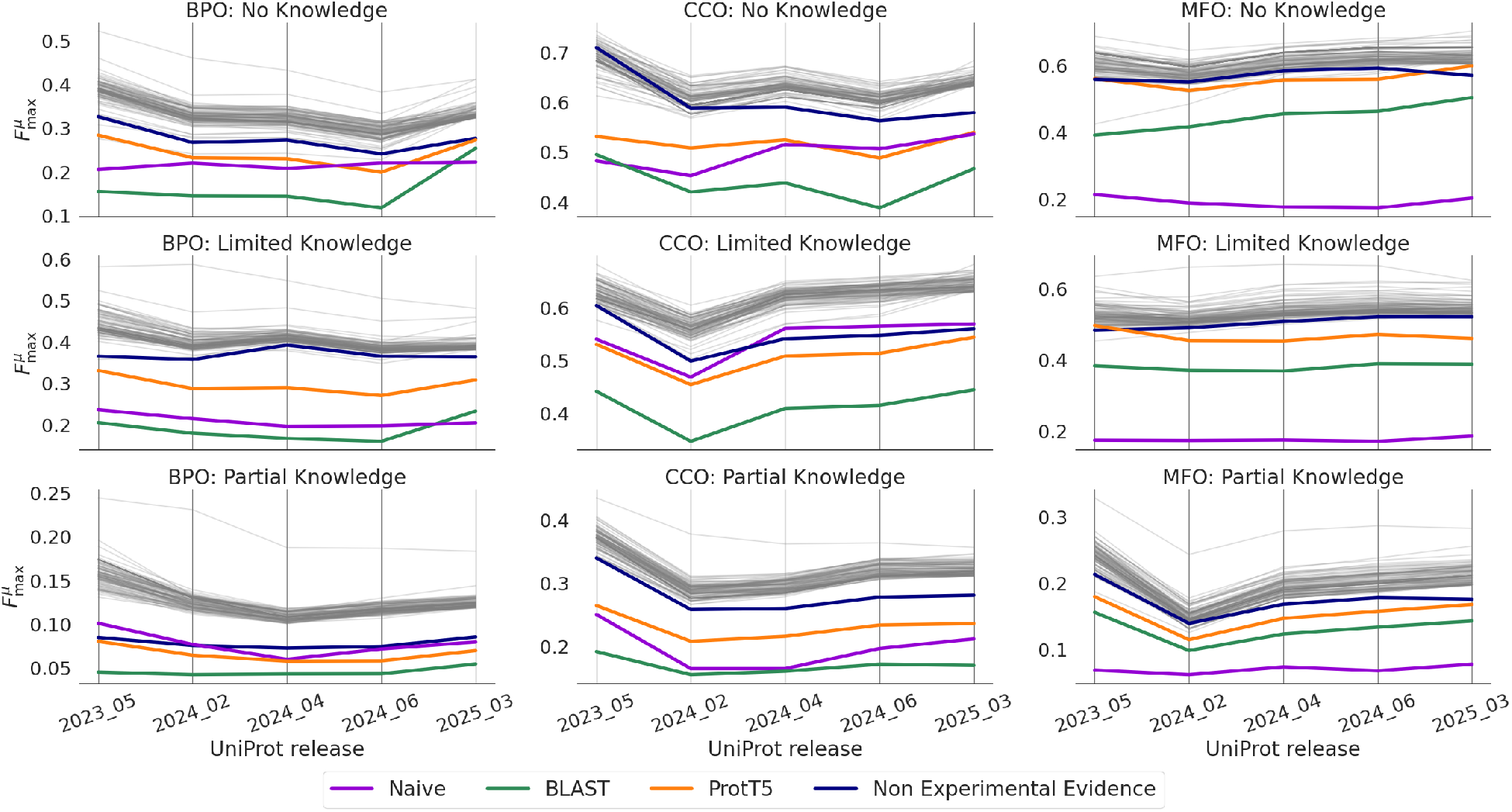
Repeated evaluation on UniProt releases since submission deadline. We plot baseline methods and top 100 submissions, as ranked by performance at the final evaluation.

## 4 Conclusion

Predicting protein function is not only a challenging modeling task, but an essential one. Proteins are responsible for undertaking nearly all physiological functions in viruses and living organisms. Understanding protein function can explain the underlying machinery of life and reveal mechanisms of disease. These insights may be key in advancing precision medicine, biotechnology, and material science [31]. Machine learning models are key to accelerating these discoveries and accurately evaluating them is crucial [13, 36]. The concerted, ongoing efforts of assessments help form and maintain communities that can come together to solve big scientific problems.

The fifth CAFA challenge, in partnership with the Kaggle platform, resulted in a substantial increase in the number of participants and submitted models. This format expanded the reach and awareness of the protein function prediction problem to new communities while still allowing domain experts to participate. Encouraging contribution from machine learning experts outside the field was one motivating factor for the change of platform. Indeed, this CAFA challenge saw a 22-fold increase in the number of participating teams, many comprised of members from outside the bioinformatics and protein fields. Increased participation, especially outside the protein prediction community, should not necessarily result in better performing models. However, the evaluation of CAFA 5 models and previous models on more than 160,000 terms showed an improvement of performance of new models. Some of this performance improvement could be attributed to more and more up-to-date training data. However, CAFA 5 models perform in general better than previous models even when the more recent BLAST and Naive baseline do not. Furthermore, the strong performance of the protein language model (pLM) embedding baseline shows how the development of protein models outside the function prediction space can help support the development of function predictors. The new pLM embedding and the non-experimentally validated annotations baselines provide a “floor” of performance and stronger benchmarks in most cases than the traditional BLAST and Naïve baselines. Model developers can use these tools as a starting point for models or to construct and supplement their training data. While we do not consider a structure-based baseline, the availability and easy usability of models such as AlphaFold2 [21] may also contribute to improvements in the function prediction field in the future.

The introduction of the partial knowledge evaluation setting allowed us to significantly expand the evaluation for CAFA5, a 229% increase in the number of annotations and 508% in the number of proteins. As more annotations accumulate in proteins with existing experimental annotations, this setting may account for an even larger proportion of function prediction evaluation. We see that model performance lags for this setting compared to the traditional settings for evaluation. This may be due to the increased difficulty of the prediction or due to lack of attention on this setting by model developers. Now that the partial-knowledge evaluation has been introduced, there is an opportunity for models to focus on this problem space. Furthermore, model developers may build models that exploit already-known annotations within a GO aspect to predict remaining functions within the same aspect. The large gap in performance compared to other settings points to the potential for improvements of this prediction task.

Finally, we validate the importance of continued evaluation across CAFA iterations by considering the performance of models across time and on the more restricted dataset of unpublished experimental evidence. We see that performance may change substantially between repeated evaluations, until possibly converging over time. When looking at unpublished annotations, we see difference in performance, especially for the BPO and MFO aspects and for all aspects in the PK settings. These fluctuations show how evaluating protein function is not a single snapshot. Benchmarks must be continuously reevaluated as the ground truth updates. What is known about protein function will always be incomplete as testing for all functions, under all contexts, for all proteins is not feasible. For this reason, computational predictors can play a significant role in scientific advancement. Yet, evaluation of these methods on the most updated scientific knowledge is necessary to gauge their utility and provide paths for improvement.

## Acknowledgments and Notices

This CAFA challenge has been partially supported by the NIH award R01GM145937 (I.F.). The contributions of the NIH author(s) are considered Works of the United States Government. The findings and conclusions presented in this paper are those of the author(s) and do not necessarily reflect the views of the NIH or the U.S. Department of Health and Human Services. Due to medical considerations S.E.B. was unable to fully review this manuscript.

## Kaggle Competition Participants

Federico Bianca ^17^, Frimpong Boadu ^18,19^, Nicola Bordin ^9^, Alberto Cabri ^20^, Elena Casiraghi ^20^, Emanuele Cavalleri ^20^, Jeroen Cerpentier ^21^, Jianlin Cheng ^18^, Alexander Chervov ^22^, Zong Ming Chua ^23^, Kerr Ding ^24^, Warith Eddine Djeddi ^25^, Tunca Dogan ^26,27,28^, Antonina Dolgorukova, Abhishek Dutta ^30^, Adibvafa Fallahpour ^31^, Sergei Fironov, Paolo Fontana ^33^, Marco Frasca ^20^, Lydia Freddolino ^34,35^, Jessica Gliozzo ^20^, Giuliano Grossi ^20^, Minh TúHoàng^36^, Robert Hoehndorf ^37,38^, Zeyu Huang^41^, Nabil Ibtehaz^42^, Emilio Ispano ^17^, Takuya Ito^44^, Joni Juvonen^45^, Dariusz Kłeczek^46^, Daisuke Kihara ^42,43^, Mesih Kilinc ^2^, Maxat Kulmanov ^37,38,39^, Enrico Lavezzo ^17^, Weining Lin ^47^, Quancheng Liu ^35^, Tong Liu^48^, Jiaqi Luo^24^, Yunan Luo ^24^, Kunal Malhotra ^49^, Mahta Mehdiabadi ^4^, Marco Mesiti ^20^, Tomi Nieminen^45^, Kexin Niu ^40^, Duc Nguyen^49^, Andrey Ogurtsov^50^, Alberto Paccanaro ^51^, Paolo Perlasca ^20^, Marina Pominova ^62^, Adarsh Rajesh ^23^, Takahiro Saito^52^, Raman Samusevich ^53,54^, Gabriela María del Mar Sánchez Rojas^51^, Ender Saribay ^26^, Stefan Stefanov ^55^, Ali Baran Tasdemir ^26^, Stefano Toppo ^17^, Mateo Torres ^51^, Erva Ulusoy ^27^, Anton Vakhrushev, Giorgio Valentini ^20^, Vlad Vinogradov ^63^, Zheng Wang ^48^, Alex Warwick Vesztrocy ^56^, Yijie Xu ^45^, Yuichiro Yoshida^57^, Atsushi Yoshizawa ^58^, Chengxin Zhang ^59^, Chenguang Zhao ^60^, Fernando Zhapa-Camacho ^37^, Shanfeng Zhu ^61^,

## Author and Competition Participant Affiliations

^1^Khoury College of Computer Sciences, Northeastern University, Boston, MA, USA ^2^Bioinformatics and Computational Biology Program, Iowa State University, Ames, IA, USA ^3^Department of Veterinary Microbiology & Preventive Medicine, Iowa State University, Ames, IA, USA ^4^Department of Biomedical Sciences, University of Padova, Padova, Italy ^5^Kaggle, Google, Mountain View, CA, USA ^6^European Molecular Biology Laboratory, European Bioinformatics Institute (EMBL-EBI), Cambridge, UK ^7^School of Computation Information and Technology (CIT), Chair for Bioinformatics, Technical University of Munich (TUM) & Institute for Advanced Study (TUM-IAS) & TUM School of Life Sciences (WZW), Munich, Germany, ^9^Institute of Structural and Molecular Biology, University College London, London, UK, ^10^Department of Ecology and Evolution, University of Lausanne, Lausanne, Switzerland, ^11^SIB Swiss Institute of Bioinformatics, Lausanne, Switzerland, ^12^Department of Biological Sciences, Carnegie Mellon University, Pittsburgh, PA, USA, ^13^Ray and Stephanie Lane Computational Biology Department, Carnegie Mellon University, Pittsburgh, PA, USA, ^14^University of California, Berkeley, CA, USA, ^15^Department of Biomedical Informatics, University of Colorado Anschutz, Aurora, CO, USA, ^16^Center for Information Technology, National Institutes of Health, Bethesda, MD, USA, ^17^Department of Molecular Medicine, University of Padova, Padova, Italy, ^18^Department of Electrical Engineering and Computer Science, University of Missouri, Columbia, MO, USA, ^19^NextGen Precision Health, University of Missouri, Columbia, MO, USA, ^20^Department of Computer Science “Giovanni degli Antoni”, Università degli Studi di Milano, Milan, Italy, ^21^Department of Electrical Engineering (ESAT), KU Leuven, Gent, East Flanders, Belgium, ^22^Unit 900, Institut Curie, Paris, France, ^23^Cancer Genome and Epigenetics Program, Sanford Burnham Prebys Medical Discovery Institute, La Jolla, CA USA ^24^School of Computational Science and Engineering, Georgia Institute of Technology, Atlanta, GA, USA, ^25^University Tunis EL Manar, Kef, Tunisia, ^26^Department of Computer Engineering, Institute of Informatics, Hacettepe University, Ankara, Turkey, ^27^Department of Bioinformatics, Graduate School of Health Sciences, Hacettepe University, Ankara, Turkey, ^28^Department of Health Informatics, Graduate School of Health Sciences, Hacettepe University, Ankara, Turkey, ^30^Department of Electrical & Computer Engineering, University of Connecticut, Storrs, CT, USA, ^31^University of Toronto, Vector Institute, University Health Network, Arc Institute, Toronto, ON, Canada, ^33^Research and Innovation Center, Edmund Mach Foundation, San Michele all’Adige, Italy, ^34^Department of Biological Chemistry, University of Michigan, Ann Arbor, MI, USA, ^35^Department of Computational Medicine and Bioinformatics, University of Michigan, Ann Arbor, MI, USA, ^36^Huê’City, Vietnam, ^37^Computer, Electrical and Mathematical Sciences and Engineering, King Abdullah University of Science and Technology, Thuwal, Saudi Arabia, ^38^KAUST Center of Excellence for Smart Health, King Abdullah University of Science and Technology, Thuwal, Saudi Arabia, ^39^KAUST Center of Excellence for Generative AI, King Abdullah University of Science and Technology, Thuwal, Saudi Arabia, ^40^Biological and Environmental Science and Engineering (BESE) Division, King Abdullah University of Science and Technology, Thuwal, Saudi Arabia, ^41^Viterbi School of Engineering, University of Southern California, Los Angeles, CA, USA, ^42^Department of Computer Science, Purdue University, West Lafayette, IN, USA, ^43^Department of Biological Sciences, Purdue University, West Lafayette, IN, USA, ^44^ITSearch Inc., Tokyo, Japan, ^45^Finland, ^46^Poland, ^47^Division of Biosciences, University College London, London, England, United Kingdom, ^48^Department of Computer Science, University of Miami, Coral Gables, FL, USA, ^49^School of Electrical Engineering and Computer Science, The Pennsylvania State University, University Park, PA, USA, ^49^Canada, ^50^Kyiv, Ukraine, ^51^Escola de Matemática Aplicada, Fundação Getúlio Vargas, Rio de Janeiro, RJ, Brazil, ^52^Tokyo, Japan, ^53^Institute of Organic Chemistry and Biochemistry of the Czech Academy of Sciences, Prague, Czechia, ^54^Czech Institute of Informatics, Robotics and Cybernetics, Czech Technical University, Prague, Czechia, ^55^Sofia, Bulgaria, ^56^BioSoft Research UK, London, England, United Kingdom, ^57^Japan, ^58^Discovery Chemistry Department; Modeling & Informatics Group, PeptiDream Inc., Kawasaki, Kanagawa, Japan, ^59^CAS Key Laboratory of Quantitative Engineering Biology, Shenzhen Institute of Synthetic Biology, Shenzhen Institutes of Advanced Technology, Chinese Academy of Sciences, Shenzhen, China, ^60^Computer and Information Sciences Department, St. Ambrose University, Davenport, IA, USA, ^61^Institute of Science and Technology for Brain-Inspired Intelligence and MOE Frontiers Center for Brain Science, Fudan University, Shanghai, China, ^62^University of Antwerp, Antwerpen, Belgium, ^63^Optic Inc., San Francisco, CA, USA

## Appendix

### A Partial Knowledge Evaluation

For partial knowledge evaluation, we must derive the expression for the marginal probability that a protein is annotated with terms (functions) in a subgraph *T*_1_ ⊂ *G* in the ontology *G*, given the *previously* annotated subgraph *T*_0_ ⊂ *G* in the same ontology for each protein. That is, we need Pr(*T*_1_|*T*_0_). This measure is used to calculate the information content of the prediction subgraph *T*_1_ conditioned on already known annotations *T*_0_. The information content is used to calculate the information-theoretic metrics misinformation, remaining uncertainty, and semantic distance [10].

#### Definitions

An ontology is represented as a directed acyclic graph *G* = (*V, E*), where *V* = {*v*_1_, …, *v*_*n*_}is the set of nodes (vertices) and *E* ⊂*V* ×*V* the set of directed edges. An edge *u* →*v* ∈ *E* is directed from node *u* to node *v* and denotes that *u* is a *parent* node of *v* and conversely that *u* is a *child* node of *v*. We say that *u* is and *ancestor* of *v* and that *v* is a *descendant* of *u* if there exists a path *u*, …, *v* in *G*. Let 𝒜 (*v*) and 𝒟 (*v*) be the set of all ancestors and the set of all descendants of node *v*, respectively. Let 𝒫 (*v*) represent the set of parents (direct ancestors) of node *v*.

In an ontology where edges encode hierarchical relationships between nodes, the *consistency requirement* states that an annotated graph is consistent if when node *v* is annotated, so are all of its ancestors, 𝒜 (*v*). For example, a protein annotated with the term GO:0042221 “response to chemical” necessarily implies it is annotated with the ancestor terms GO:0050896 “response to stimulus” and GO:0008150 “biological process” through the hierarchical *is a* relation encoded by the edges between these term nodes.

#### Information theoretic metrics for partial knowledge evaluation

Let *x* ∈ *𝒳* be inputs with labels *Y* ∈ *𝒴* that are consistent directed acyclic graphs describing annotations in an ontology. Assume that the ontology label distribution can be factorizes as a Bayesian network as in [10]. That is, the marginal probability of a target *x* ∈ 𝒳 being annotated with terms associated with nodes *{v} ∈ T ⊆ G* is given by

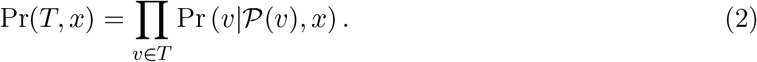

##### Lemma 1.

*Assume an ontology graph G has label annotations generated according to the probability distribution from a Bayesian network following the ontology structure. For an annotated consistent subgraph T*_0_ ⊆*G, and new annotations on nodes* {*v*} ∈ *T*_1_ \*T*_0_ ⊆*G, the information content of the new annotations is*

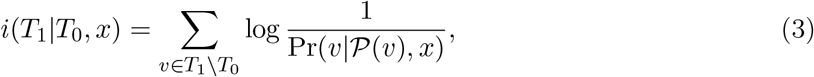

*where T*_1_ *\T*_0_ *is the subgraph induced by the set of nodes that are in subgraph T*_1_ *but are not in subgraph T*_0_.

*Proof*. For notational convenience, we drop the implicit dependence on *x*. Assume no annotations were removed so that at time *t*_1_, a target *x* is associated with the annotated terms at *t*_0_ and those added at *t*_1_, i. e. the nodes {*v*} ∈ *T*_1_ *∪ T*_0_ *⊆ G*. Then, because *T*_1_ is a consistent subgraph of *G, T*_0_ *\T*_1_ = ∅ and *T*_0_ *∩ T*_1_ = *T*_0_. Thus, the conditional probability becomes

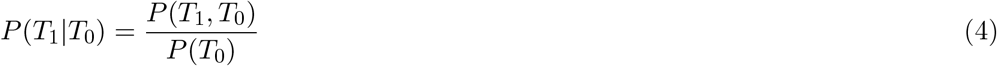

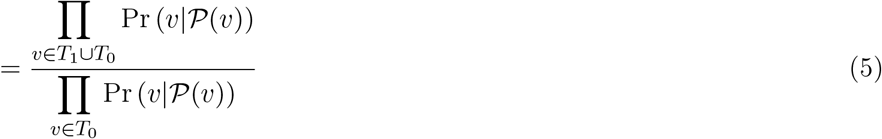

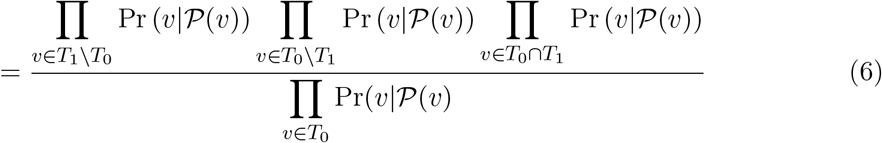

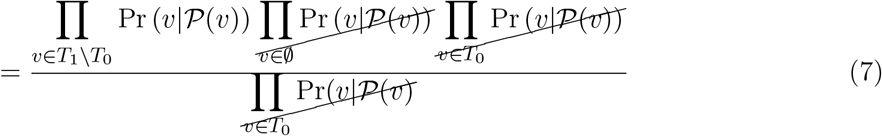

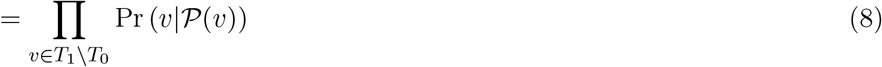

Substituting the above into the definition of information content,

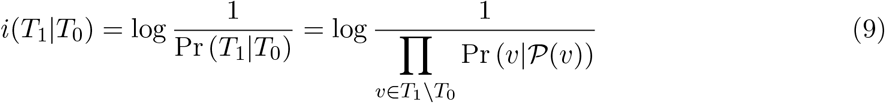

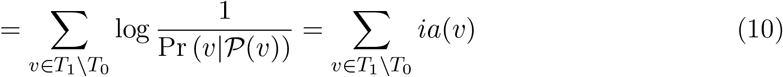

That is, the information content of the consistent subgraph induced by newly annotated nodes is equal to the information content *only* from newly annotated nodes.

### B Implementation Details

#### B.1 Evaluation

We adapt the CAFA-evaluator [26] code base to account for PK evaluation and share the code here: https://github.com/claradepaolis/CAFA-evaluator-PK. To evaluate predictions, we run the code as follows 

~~~
cafaeval [onotology file] [prediction directory] [gt terms] -ia [ia file] -toi [terms of interst] -prop fill -th_step 0.001 -no_orphans
~~~

For partial-knowledge evaluation,

~~~
cafaeval [ontology file] [prediction directory] [gt terms] -ia [ia file] -known [terms to exclude] -toi [terms of interst] -prop fill -th_step 0.001 -no_orphans
~~~

#### B.2 Data Versions and Sources

For ground truth annotations, we use annotations with GO evidence codes as listed in Table A1. Annotations with these evidence codes are also used to construct the known annotations at *t*_0_ to be excluded when evaluating the partial knowledge setting.

Details of each dataset used in evaluations is listed in Table A2. The release and data for each UniProt, GOA, and the GO ontology are listed for each time point. The corresponding time points for previous CAFA challenges is also listed.

**Table A1:**
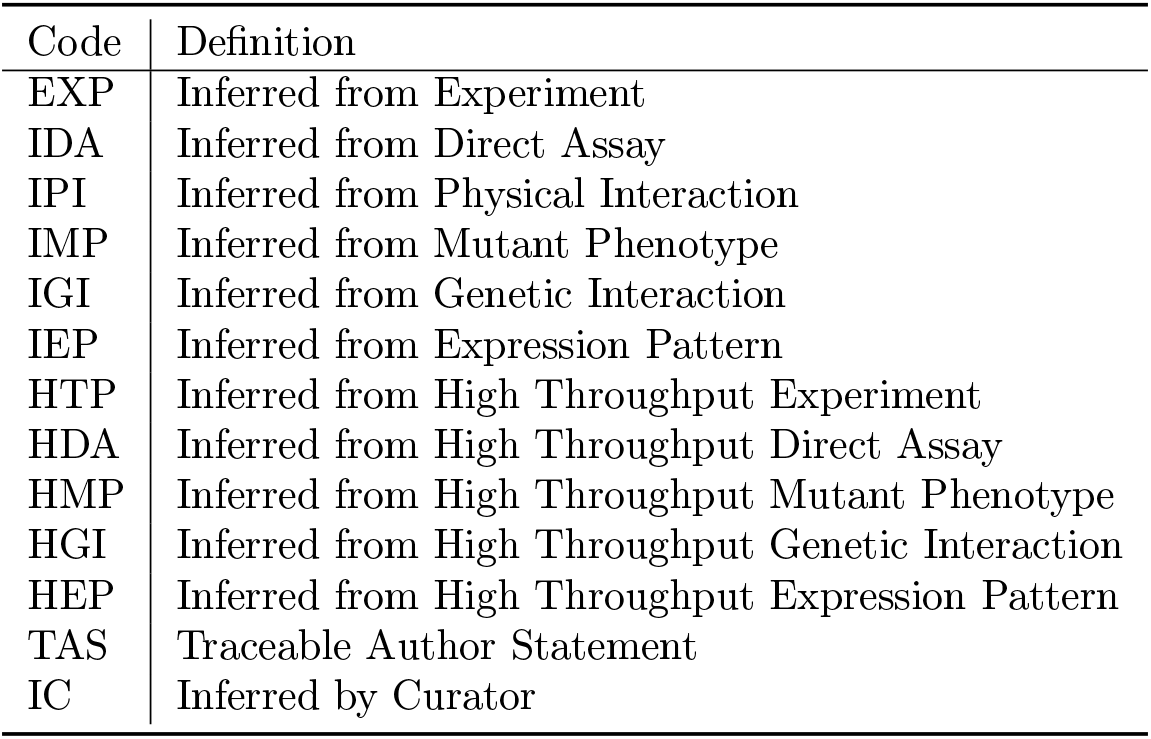
Evidence codes used in CAFA 5 evaluation and to filter out previously-known annotations.

**Table A2:** Data time points and version for each evaluation.

| Time | UniProt |  | Gene Ontology |  | Use |
| --- | --- | --- | --- | --- | --- |
|  | Version | Date | GOA | GO |  |
| $t_\ell$ | 2022_05 | | | 2023-01-01 | Target protein and terms frozen |
| $t_0$ | 2023_03 | 2023-08-21 | 2023-07-12 | 2023-07-27 | Filter out all known annotations at this date;<br>Used for **BLAST** and **Emb** baselines look-up |
| $t_e$ | 2023_05 | 2023-11-08 | 2023-12-04 | 2023-11-15 | Kaggle competition results; intermediate evaluation |
|  | 2024_02 | 2024-03-27 | 2024-04-16 | 2024-04-24 | Intermediate evaluation |
|  | 2024_04 | 2024-07-24 | 2024-08-01 | 2024-06-17 | Intermediate evaluation |
|  | 2024_06 | 2024-11-27 | 2024-12-21 | 2024-11-03 | Intermediate evaluation |
|  | 2025_03 | 2025-06-18 | 2025-05-03 | 2025-06-01 | Final evaluation |
| Head-to-head CAFA Evaluation |  |  |  |  |  |
| $t_\ell^{\text{CAFA2}}$ | | Sep 2013 | | 2013-08-01 | GO terms predicted in CAFA 2 |
| $t_0^{\text{CAFA2}}$ | 2013_01 | Jan 2014 | 2013-01-07 | | Used for head-to-head **BLAST** baselines look-up for CAFA 2 |
| $t_\ell^{\text{CAFA3}}$ | | Sept 2016 | | 2016-08-01 | GO terms predicted in CAFA 3 |
| $t_0^{\text{CAFA3}}$ | 2017_01 | Feb 2017 | 2017-01-17 | | Used for head-to-head **BLAST** baselines look-up for CAFA 3 |
| $t_\ell^{\text{CAFA4}}$ | | 2019-10-16 | | 2019-10-07 | GO terms predicted in CAFA 4 |
| $t_0^{\text{CAFA4}}$ | 2019_01 | 2020-02-12 | 2019-12-17 | | Used for head-to-head **BLAST** baselines look-up for CAFA 4 |

### C Additional Results

**Figure A1:**
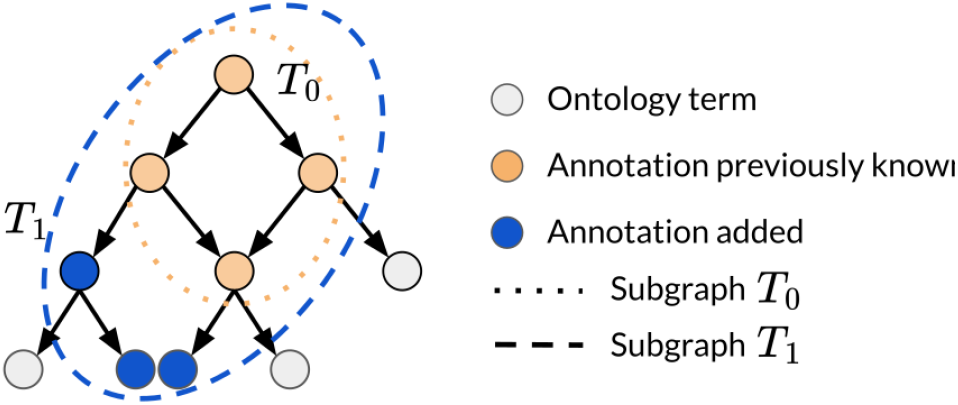
Known annotations (yellow) in subgraph *T*_0_ and new annotations (blue) added. Only new annotations *T*_1_ *\ T*_0_ are evaluated in partial knowledge evaluation.

**Figure A2:**
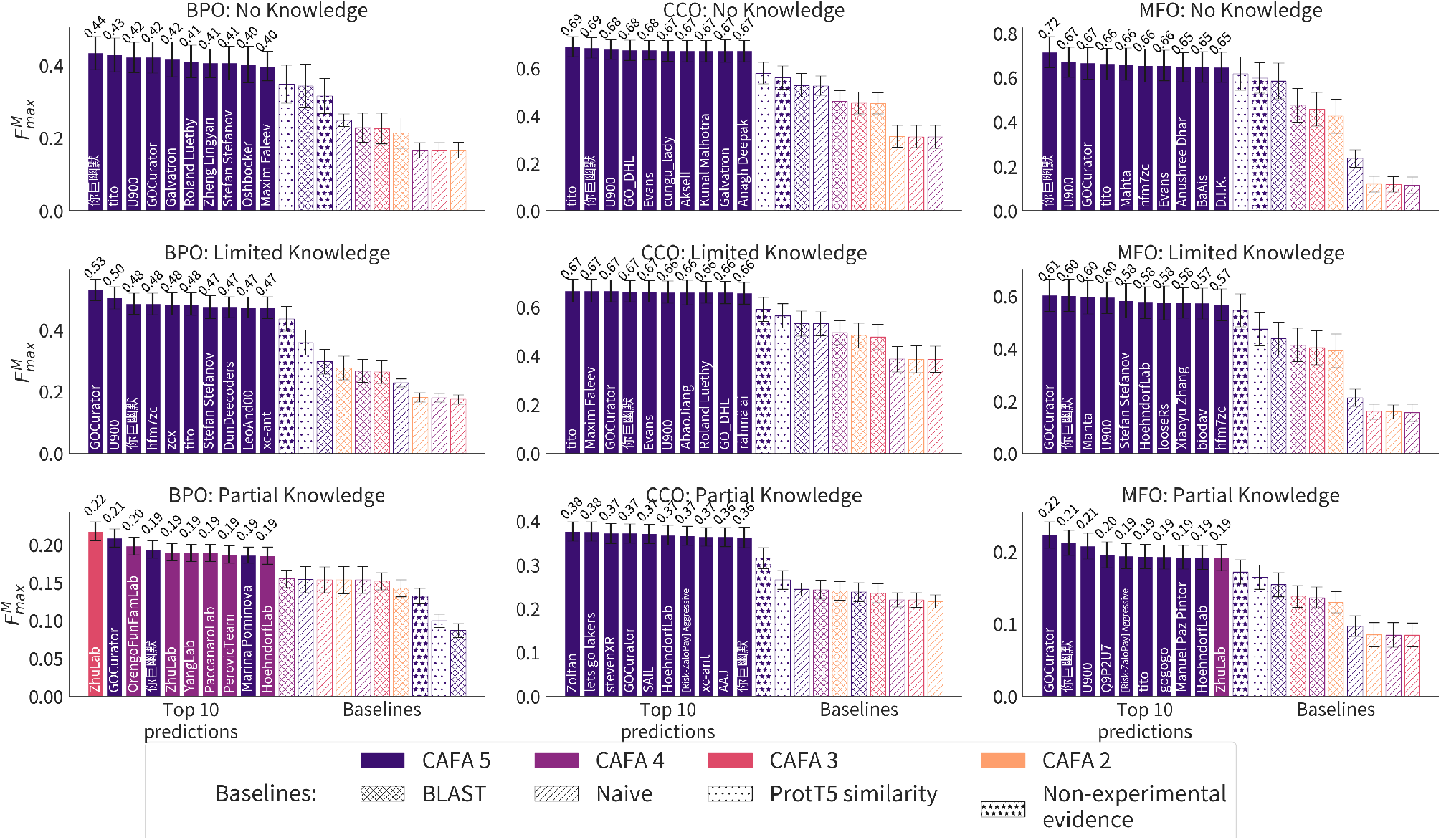
Macro-average *F*_max_ for 5,000 bootstrap samples with 95% confidence interval error bars. Group names are shown in bars. BPO: Partial Knowledge is the only metric and setting with multiple teams from previous CAFA challenges in the top-10 predictors

**Figure A3:**
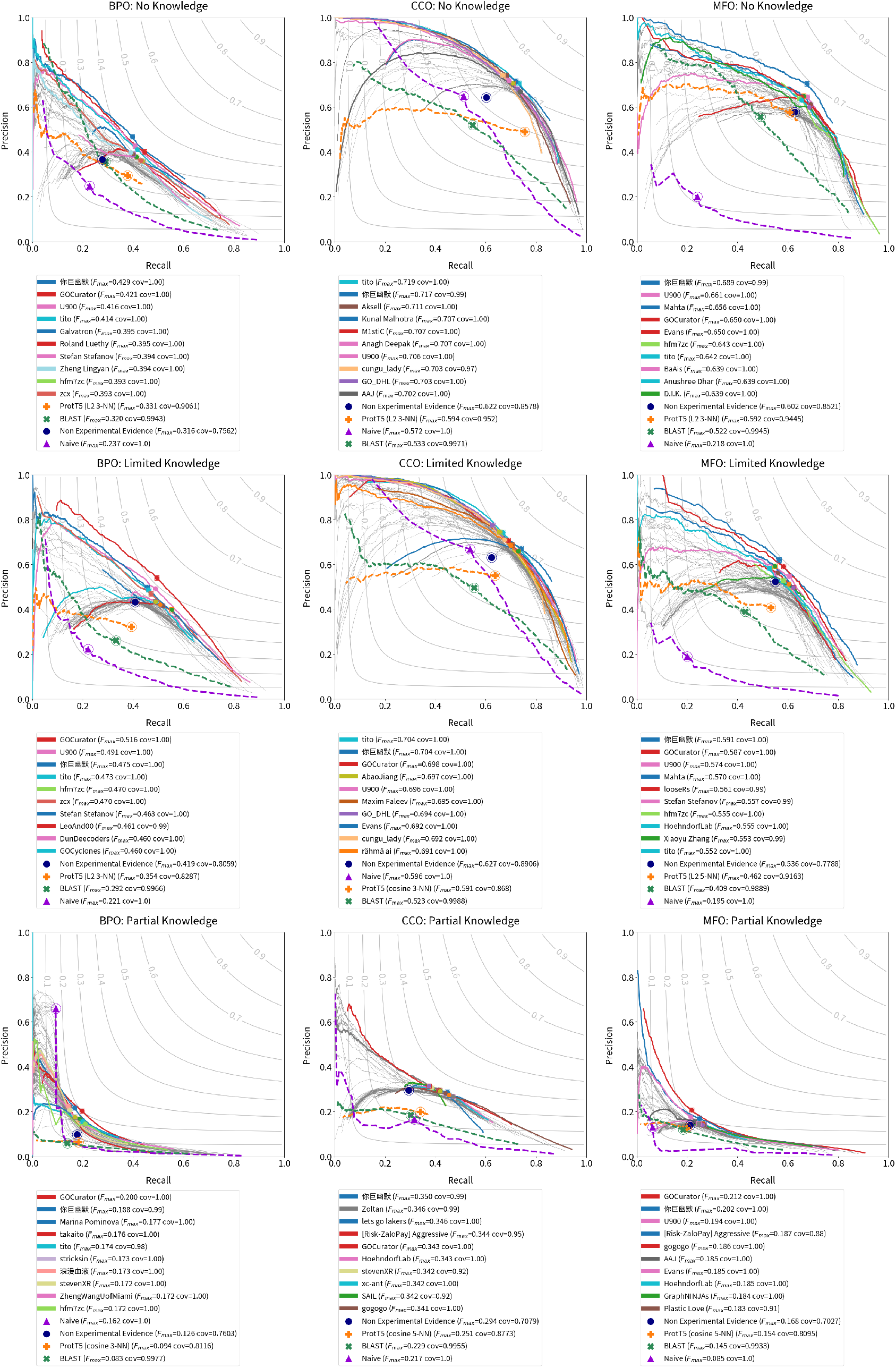
Macro-average precision/recall curves for top CAFA 5 models.

**Figure A4:**
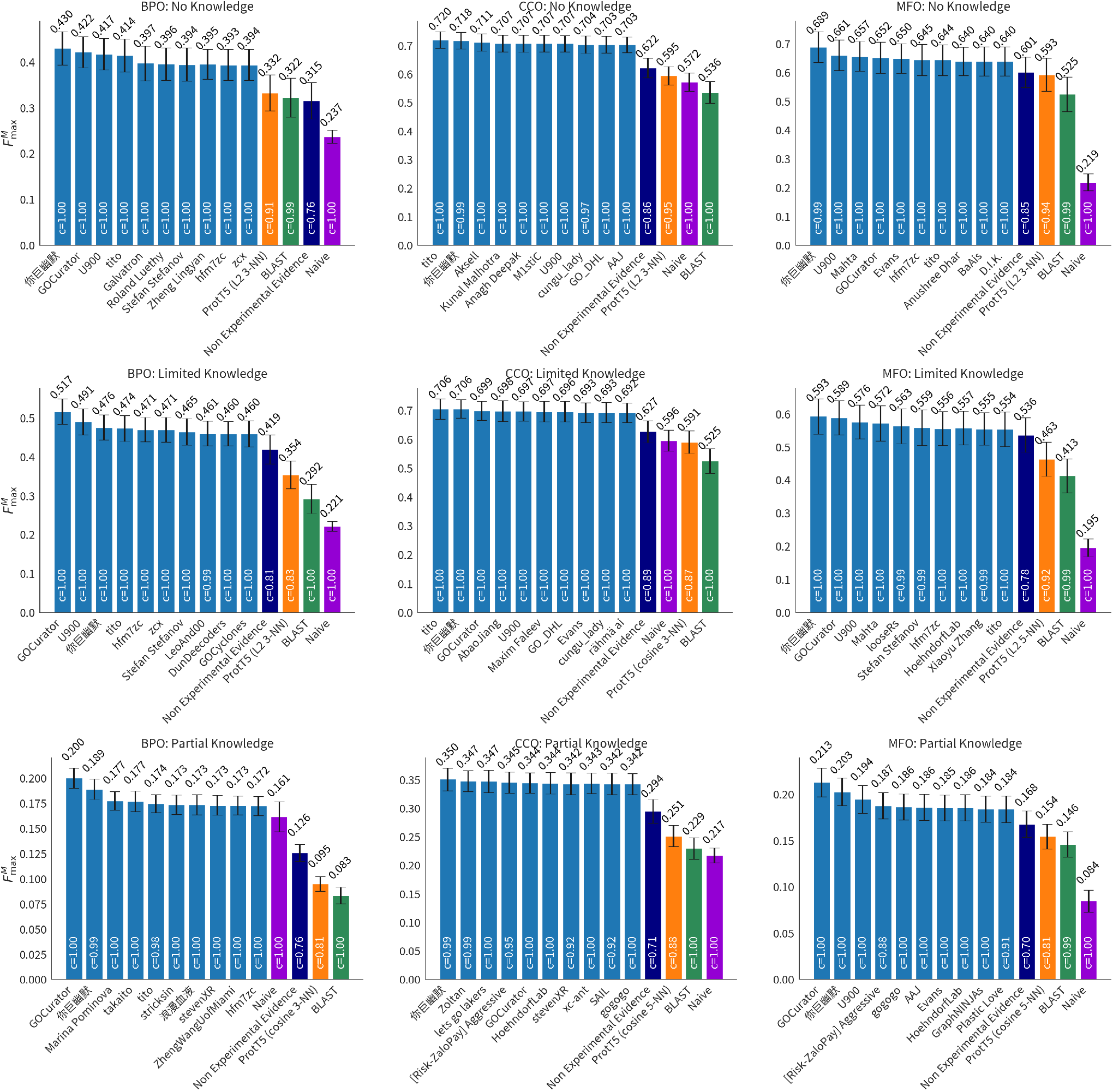
Macro-average 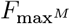 for 5,000 bootstrap samples for top CAFA 5 models.

**Figure A5:**
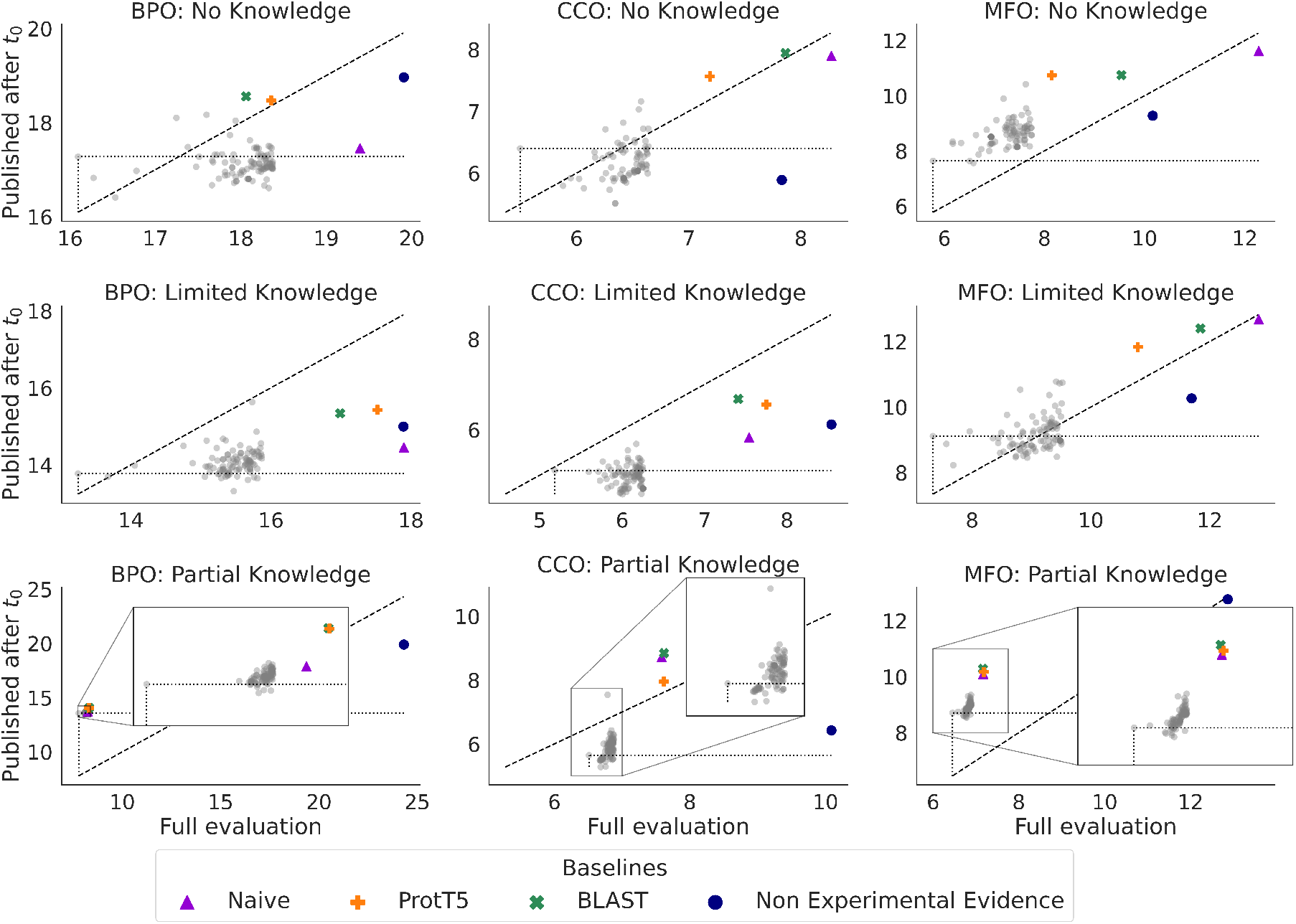
We compare *S*_min_ of methods on the full evaluation set (horizontal axis) with a *S*_min_ on just the subset of annotations from publications released after *t*_0_ (vertical axis). Baselines and the best 100 methods are shown. Points below the diagonal line indicate that the method’s *S*_min_ improved when evaluated on the subset of annotations from new publication. Points above the diagonal line indicate an increase in *S*_min_. The *S*_min_ of the top-performing method in the full evaluation is designated by a dotted line. In each case, the top-performing model on the full evaluation is not the best-performing when evaluated on newly-published annotations. That is, there are other points *below* the horizontal dotted line.

**Figure A6:**
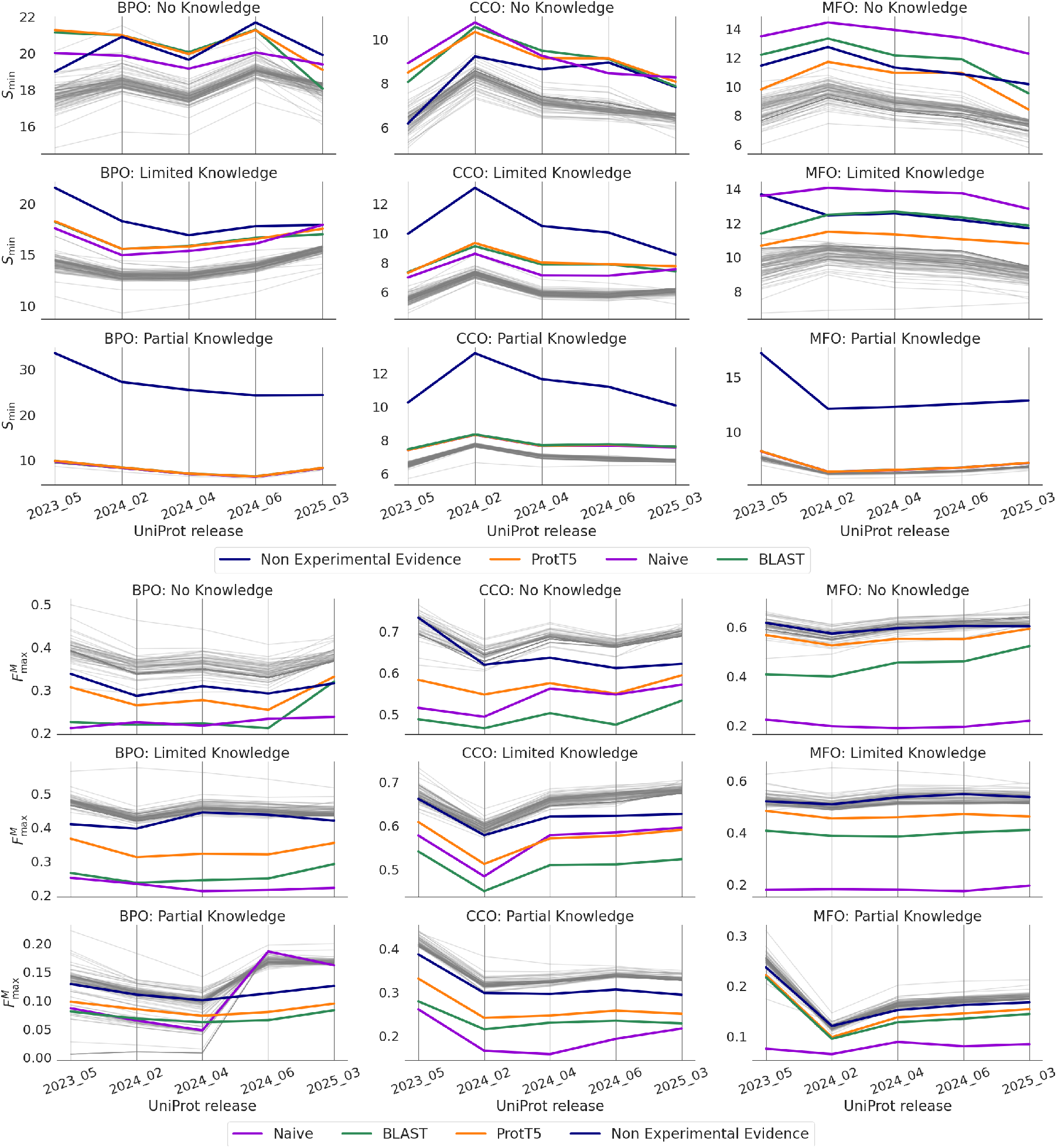
Performance of top 100 models with respect to *S*_min_ (top) and 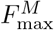 (bottom) for UniProt releases over time.

## Footnotes

1 Evaluation code is available at https://github.com/claradepaolis/CAFA-evaluator-PK/. Baselines and evaluation data are available at https://doi.org/10.5281/zenodo.20186533.

